# Nearly (?) sterile avian egg in a passerine bird

**DOI:** 10.1101/2023.02.14.528324

**Authors:** Martin Těšický, Lucie Schmiedová, Tereza Krajzingrová, Mercedes Gomez Samblas, Petra Bauerová, Jakub Kreisinger, Michal Vinkler

**Author notes:** <u>Author for correspondence:</u> Martin Těšický, Correspondence address: Martin Těšický, Charles University, Faculty of Science, Department of Zoology, Viničná 7, 128 43 Prague, Czech Republic, EU. **Author’s Contribution** M.T., M.V. and J.K. designed the study. M.T., T.K., P.B. and M.V. collected samples. M.T., L.S., and T.K. with help of M.G. performed the laboratory analysis. M.T. and L.S. with help of J.K. analysed data. M.T. and M.V. interpreted the data and drafted the manuscript. All authors contributed their comments to the final approved version of the manuscript. **Ethical approval** The research was carried out under the current laws of the Czech Republic and the European Union. The experiment was approved by the Environmental Protection Department of Prague City Hall (permit No. S-MHMP-1061728/2010/OPP-V-790/R-235/Bu) and the Ministry of the Environment of the Czech Republic (permit No. 22003/ENV/16-1009/630/16). Permission for catching and ringing adult birds was granted by the Bird Ringing Centre of the National Museum in Prague.

## Abstract

During early ontogeny, microbiome affects development of the gastrointestinal tract, immunity and survival in vertebrates. Bird egg has been suggested to be either (1) initially sterile (*Sterile egg hypothesis*) and (2) colonized through *horizontal trans-shell migration* after egg laying, or (3) initially seeded with bacteria through *vertical transfer* from mother’
ss oviduct. Little empirical data illuminate so far the contribution of these mechanisms to gut microbiota formation in avian embryos. We evaluated microbiome of the egg content (day 0; E0-egg), embryonic gut at day 13 (E13) and female faeces in a free-living passerine, the great tit (*Parus major*), using a methodologically advanced procedure combining *16S rRNA* gene sequencing and microbe-specific qPCR assays. Our metabarcoding revealed that avian egg is (nearly) *sterile*, barely distinguishable in microbial composition from negative controls. Of the three potentially pathogenic bacteria targeted by qPCR, only *Dietzia* was found in E0-egg (yet also in controls), E13 gut and female samples, which might indicate its possible *vertical transfer*. Unlike in poultry, we have shown in passerines that major bacterial colonisation of the gut does not occur before chick’s hatching. We stress that protocols carefully checking for environmental contamination are critically important in studies using samples with low bacterial biomass.

## Introduction

Gut microbiota plays a paramount role in host physiology, affecting nutrient digestion (Bäckhed *et al*. 2005), gastrointestinal tract (GIT) and immune system regulation (Ost and Round 2018), gut-brain-axis signalling (Strandwitz 2018), and also the onset of diseases (Honda and Littman 2012). While early microbial next-generation sequencing (NGS) studies suggested that transmission of the maternal microbiome before birth may be a universal phenomenon in both vertebrates and vertebrates, including humans (Funkhouser and Bordenstein 2013), some earlier studies are now thought to have suffered from the increased rates of environmental contaminants and sequencing artefacts as metabarcoding protocols have advanced. In humans, the current consensus is that bacteria are not present in placenta and foetus under physiological conditions but they colonize new-born gut since birth (de Goffau *et al*. 2019) but see Walker *et al*. (2017). Is a bird egg also initially sterile or do females deposit any bacteria into the egg to direct the initial embryonic and chick microbiome development? Although with the advent of NGS, our understanding of avian microbiota composition exponentially increased over the last decade (Sun *et al*. 2022), this improvement concerns mostly avian adults, with little being known about the bacterial colonization of the bird egg and microbial communities formed during embryonic development. Early bacterial colonisers may shape GIT and the immune system of developing embryos and also influence their survival and the composition of chick microbial communities after hatching (Hansen *et al*. 2015; Roto, Kwon and Ricke 2016). However, based on a standard search through Web of Science (5^th^ of September 2022), there were 554 papers dealing with microbiota in birds; with the following keywords searched in topics: (microb* OR bacter*) AND (bird* OR avian*) AND 16S rRNA, but only 22 papers dealing with microbiota in egg content or embryo [(microb* OR bacter*) AND (bird* OR avian*) AND 16S rRNA AND (egg white OR albumen* OR egg content OR embryo*)], out of which only four articles were relevant to the topic of the study.

*Sterile egg hypothesis* assumes that bird egg is initially formed sterile in the female reproductive tract. i.e. bacteria are not maternally vertically transmitted to the egg content in this developmental phase (Roto, Kwon and Ricke 2016). This would require either the absence of bacteria in the oviduct tract or, if this is not the case, presence of host physiological filters in oviduct tissue that prevent bacteria from colonising the developing egg (Lee *et al*. 2019). In this scenario, bacteria occurring later on in the eggs would need to colonize the eggs during laying and incubation period via their eggshell pores (Bruce and Drysdale 1994). The *sterile egg hypothesis* was particularly influential in the era of cultivation-based studies (reviewed in Roto, Kwon and Ricke 2016) but recently some egg microbial communities have been detected by 16S rRNA gene metabarcoding (see below). Bird eggs are well protected by several physical (four demarking layers, i.e. cuticle covering also eggshell pore openings, crystalline eggshell and inner and outer shell membranes; Lunam and Ruiz 2000; Liong, Frank and Bailey 1997; D’Alba and Shawkey 2015) and chemical protective mechanisms (e.g. antimicrobial peptides with bacteriolytic activity secreted mainly in dense albumen, such as lysozyme, gallin, β-defensins, ovalbumin, ovotransferrin and ovocalyxin; Mann 2007; Gantois *et al*. 2009; Cuperus *et al*. 2013) from bacterial colonization. This creates a very unhostile environment for microbes and protects the germinal disc and nutrient-rich yolk. Besides that, birds can actively regulate eggshell microbiota, e.g. by incubation which may reduce overall bacteria diversity and the growth of specific pathogenic bacteria (Cook *et al*. 2005) or by coating eggshell with bactericidal oils from their uropygial glands (Soler *et al*. 2016). Despite such protection, bacteria can still colonize avian eggs via two different mechanisms (Pedroso 2009; Roto, Kwon and Ricke 2016): (1) as a *vertical transfer* of microbiota from mother oviduct to the developing egg and/or (2) as *a horizontal trans-shell migration* from the environment after egg laying.

In *vertical transfer*, it is usually assumed that the bacteria colonise the female reproductive tract, the oviduct. Live bacteria can directly access the oviduct through ascending infection from the cloaca. Alternatively, the host could support the bacterial uptake in the gut by macrophages or dendritic cells which then migrate and transport selected bacteria to the oviduct (Gantois *et al*. 2009). Certain bacteria can also have specific adaptations to increase their probability of successful egg colonisation (Song *et al*. 2022). *Vertical transfer* of pathogenic bacteria, such as *Salmonella* has been suggested in culture-based studies where experimentally orally infected hens laid contaminated eggs (Keller *et al*. 1995; Gantois *et al*. 2009; Pedroso 2009). However, the frequency of such vertical transmission is typically very low (e.g. for *Salmonella* usually ranging between 0-4.5 % in egg whites but can occasionally reach up to 20%), and significantly differs among studies, with differences between different *Salmonella* serovars and egg compartments infected (e.g. stronger contamination occurs in the egg white than in the yolk; reviewed in Gantois *et al*. 2009). Recently, an NGS-based study in Korean chickens (Lee *et al*. 2019) has revealed a diversified bacterial community in egg white and embryonic samples. Only 3.2 % bacterial taxa have been shared with oviduct microbiome, suggesting low *vertical transfer* from the reproductive tract (Lee *et al*. 2019). In another study, four passerine species had distinct microbial communities in egg whites (E0) from negative controls (Trevelline *et al*. 2018). Yet, the supporting dataset was in this study very small, consisting of only 11 eggs in total. Therefore, such evidence for the *vertical transfer* should be interpreted with caution. Furthermore, the analytical approaches used in these two studies may not reliably exclude all false positives due to contamination noise.

*Bacterial horizontal trans-shell migration* assumes that bacteria colonize avian eggs during the post-laying period (Bruce and Drysdale 1994). Originating from the nest environment (female or nest microbiome, generally termed nidobiome; Campos-Cerda and Bohannan 2020), bacteria can migrate via pores on the eggshells to reach the egg content and establish microbial communities in the embryos (e.g. Van Veelen, Salles and Tieleman 2018; Lee *et al*. 2019). Despite egg protection (see above), some bacteria, such as *Neisseria* in the Greater white-fronted geese (*Anser albifrons*; Hansen *et al*. 2015) have been documented to penetrate the eggshell, causing premature embryo mortality. Although it is difficult to distinguish *trans-shell migration* from the *vertical transfer* only from observational data, it seems that bacteria are recruited predominantly to the embryonic gut by *trans-shell migration*. This is viewed in chicken embryos where diversified microbial communities were revealed in embryonic gut both based on classical microscopy (Kizerwetter-Świda and Binek 2008) as well as more recently by 16S rRNA gene metabarcoding (e.g. Ding *et al*. 2017; Lee *et al*. 2019; Ding *et al*. 2022). Chicken embryonic microbiome is dominated by taxa, such as *Pseudomonas, Janthinobacterium, Acinetobacter, Stenotrophomonas* and *Meganomonas* (Lee *et al*. 2019). It is likely that these bacteria primarily originate from eggshell, gastrointestinal tract or cloaca of females as indicated by sharing some taxa between embryos and these compartments. For instance, 42 % of bacterial taxa from embryonic GIT have been also found in hen faeces (Ding *et al*. 2017) and in another chicken study, as much as 63.4 % and 31.7 % of the microbial taxa from the egg white and embryonic gut were found on eggshell and female cloaca, respectively (Lee *et al*. 2019). Despite the sharing of some taxa, embryonic intestine microbiome in chickens appears to be distinct from the female faeces and urogenital tract microbiome (Lee *et al*. 2019), supporting the role of some host physiological filters.

The level of bacteria colonization may vary between species, with differences e.g. between free-living and captive species or differently developed chicks (altricial vs precocial species). However, some caution is needed when interpreting the results of different NGS studies. For example, not all NGS studies adequately integrated negative controls of various types (extraction, amplification, etc.) or performed amplification in PCR replicates. Unlike the studies in chickens (e.g. Ding *et al*. 2017; Lee *et al*. 2019; Ding *et al*. 2022), Grond *et al*. (2017) revealed only negligible microbiota (not significantly different from negative controls) in embryonic GIT of two arctic shorebirds. This raises the question of whether the observed marked differences in microbiota composition of egg content and embryonic GIT in birds are due to differences between species or whether the results could also be biassed by methodological differences (such as the lack of negative controls or no verification of sequencing-based metabarcoding data by more sensitive qPCR). Therefore, the extent to which these two different, yet mutually non-exclusive mechanisms (*vertical transfer* and *horizontal trans-shell migration*) contribute to the establishment of the embryonic microbiota in wild birds is still not sufficiently understood.

In our study, for the first time in wild birds, we simultaneously assessed whether the bird egg is initially sterile (i.e. testing the *Sterille egg hypothesis*) and how bacterial colonization occurs (if any) during embryonic development (*vertical transfer* vs. *horizontal bacterial trans-shell migration*). We came up with an innovative approach to study the microbial profiles of egg content and embryonic GIT in the great tit (*Parus major*). During the breeding season of 2018, we collected a total of 240 microbial samples from 57 nests of a free-living great tit population breeding in Prague, Czech Republic. We adopted a methodologically improved 16S sRNA metabarcoding approach in combination with specific qPCR assays to reveal sample contaminants masking the natural variation in microbial composition. Our study aimed to test (i) whether the egg is initially sterile in the great tit by describing the initial egg-content microbiota sampled shortly after laying (embryonic day 0, E0) and (ii) assess the roles of different colonization mechanisms based on the comparison of the initial egg-content microbiota, embryonic GIT microbiota present shortly before hatching (embryonic day 13, E13), and female faecal microbiota. Particular attention was paid to pathogenic bacteria with potential negative effects on host fitness. We also administrated E0 eggs (*in ovo*) with *Enterococcus faecium* in two different concentrations to validate our ability to detect bacteria later in E13.

## Materials and methods

### Sampling design and sample collection

The sampling was performed in a free-living great tit population breeding in artificial nest boxes in a deciduous forest at the edge of Prague, Czech Republic, EU (50°08’12.4”N, 14°27’57.2”E; see Těšický *et al*. 2021, 2022 for more details about study site) during their breeding period on April and May 2018. In total, we collected 52 eggs to sample microbiome of egg content (E0-egg), 118 eggs to sample E13 embryonic intestine (unmanipulated, E13-nat, as well as *Enterococcus*-manipulated eggs, E13-Ent, see below), and 34 female faecal samples (see Table S2, SI1 for number of samples in different categories). In total, these samples represented 58 nests, out of which the complete sample set was available from 18 nests. The time of breeding was determined by the regular inspections of nest boxes (about 2-7 day interval, adjusted to the estimated hatching date). We numbered all eggs in the clutches based on their laying order with a permanent marker.

To describe natural microbiota composition in E0-eggs, we collected one freshly laid egg per clutch (N = 52) within one day from laying and transported it into the laboratory. Here, in a laminar biosafety cabinet (Jouan MSC 12, ThermoFisher Scientific, Carlsbad, USA) its surface was cleaned with 96 % ethanol and DNA remover to prevent contamination. Subsequently, all egg content, i.e. approximately 200–300 μL was aseptically aspirated with the insulin syringe (B Braun, Cat. No. 9151125, Melsungen, Germany) after punctuating the eggshell. Then samples were stored frozen in PCR-clean cryotubes (Simport, Canada) at −80 °C.

To describe natural microbiota composition in embryonic GIT (E13-nat), we collected either one or two E13 eggs per clutch (N = 66). Two E13 eggs per nest were collected from a total of 25 clutches to assess whether microbiota is more similar within than between nests. To confirm that putative bacteria present in E0-egg can be detected with our methods later in the life of the embryo, we injected within two days after laying a subset of the eggs (one egg per nest, N = 56) with *Enterococus faecium* (ref. strain: NCIB 11181; probiotics Lactiferm Basic 5, Chr. Hansen, Hørsholm, Denmark Cat. No. L-0265). Specifically, we injected the first egg per clutch with 10 µl of *E. faecium* in a concentration of either 10^7^ (high dose) or 10^4^ (low dose) colony-forming units (CFU; since equal results were obtained categorised jointly as E13-Ent, see the Results) and a second egg with the 10 µl of PBS (Sigma Aldrich, Cat. No. D5652-50L; E13-PBS, N = 53), serving as a control. After the manipulation, the treated eggs were sealed at the injection site with a superglue, returned to their nests and together with E13-nat were let to incubate until E13 when all E13 eggs were collected. (please, see SMMO 3 for details on the *in ovo* application, Figure 1 for a timeline and sample design scheme and Figure 2 for the method overview). The collected E13 eggs were transported into laboratory while kept in an incubator (Brinsea Octagon 20 Advance Incubator, Brinsea Products Inc, Titusville, USA) at constant temperature and humidity (37.5°C and 60 %) until their aseptic dissection (maximum 6 hours after collection). In the biosafety cabinet, embryos were aseptically removed from the eggs, and placed on sterile Petri dishes (ThermoFisher Scientific, Cat. No. 101IRR). Then embryos were decapitated and their lower gastrointestinal tracts (the part between gizzard and cloaca) were taken and stored frozen at −80 °C in RNA later (Quaigen, Cat. No. 76106, Hilden, Germany). Tissue samples were successfully obtained in a total of 23 E13-Ent and 29 E13-PBS eggs, i.e. 41.1 % and 54.7 % of initial numbers because of embryos’
s mortality.

**Figure 1:**
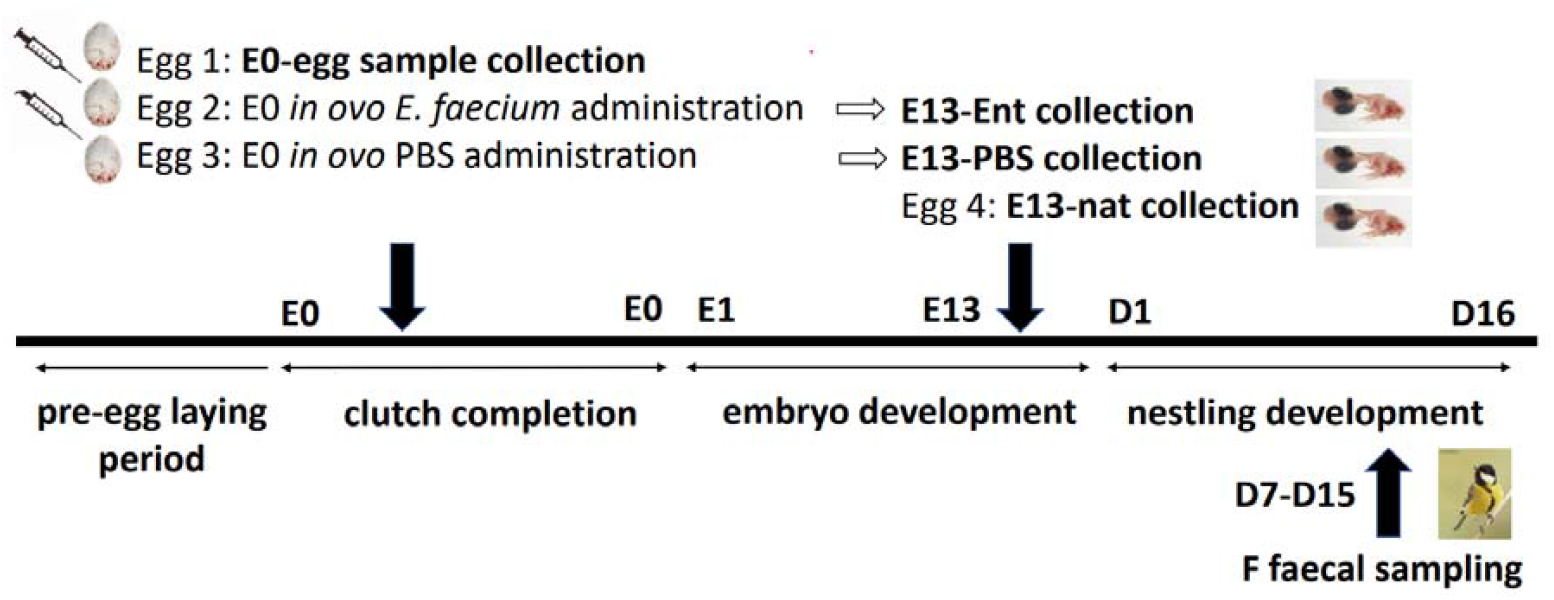
Timeline and experimental design of the great tit study. Typically, five microbiota samples per nest were collected: E0-egg – egg content sampled on embryonic day 0 (E0), E13-nat – E13 intestinal sample from a non-manipulated egg, E13-Ent – E13 intestinal sample from an *Enterococcus*-treated egg, E13-PBS – E13 intestinal sample from a control, PBS-injected egg, and F – female faecal samples collected between day 7 and day 15 (D1-D15) of the nestling age. E and D above the axis indicate the day of embryonic and chick development, respectively.

**Figure 2:**
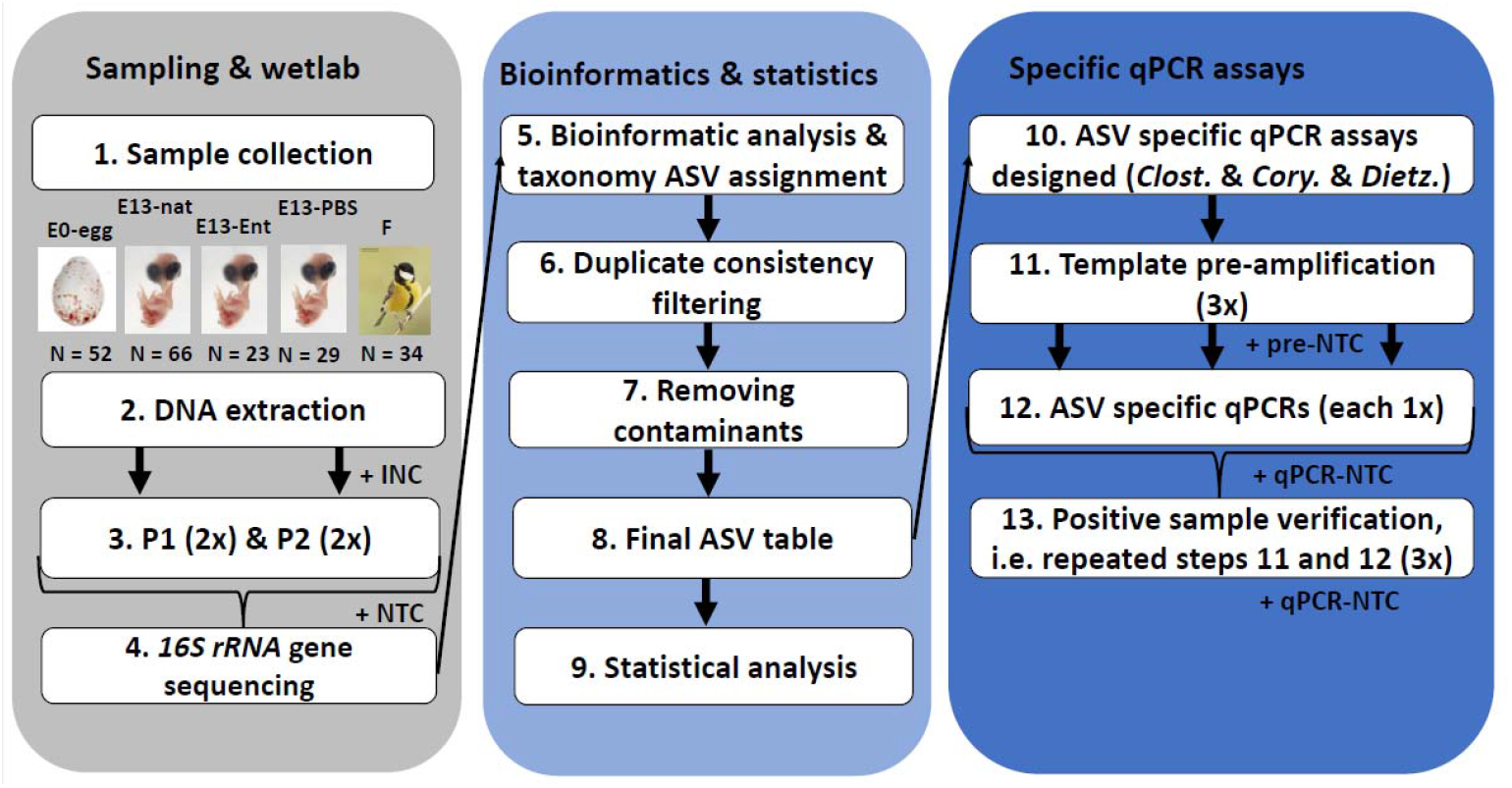
Schematic representation of the procedure used for assessing bacteria composition in egg content, embryonic intestine, and female faecal samples in the great tit. (1) In the field, different biological sample types were collected: E0-egg – egg content sample of embryonic day 0, E13-nat – E13 intestinal sample from non-manipulated egg, E13-Ent – E13 intestinal sample from *Enterococcus*-treated egg, E13-PBS – E13 intestinal sample from a control PBS-injected egg, F – adult female faecal sample. (2) Bacterial DNA was extracted including isolation negative controls (INC). We applied (3) two protocols for 16S rRNA gene DNA microbial genotyping: protocol 1, P1 with no-chloroplast amplifying primers and protocol 2, P2 with bacterial universal primers and 16S rRNA amplicon libraries were prepared. The libraries were sequenced (4) and then (5) bioinformatic analysis and taxonomical assignment of bacteria were performed. (6) Only amplicon sequence variants (ASVs) consistently present in both technical duplicates were retained. (7) We removed potential contaminants and (8) based on the abundance of particular ASVs in different biological sample types, we further (9) statistically analysed sequencing data. Based on these results, (10) we designed specific probe-based qPCR assays to detect potentially pathogenic bacteria (*Clostridium, Dietzia*, and *Corynebacterium*). (11) To increase the sensitivity of qPCR assays, we pre-amplified the template DNA with bacterial universal primers including negative control of pre-amplification (pre-NTC). (12) ASV-specific qPCRs were then performed with these pre-amplified DNA templates and negative control of qPCR (qPCR-NTC). (13) Positive samples from qPCR were further verified by independent amplification (repeated steps 11 and 12). Numbers in brackets correspond with the number of technical replicates per sample.

To describe maternal female microbiota, we also non-invasively collected faecal samples from adult females (N = 34) using our previously described methodology (Kropáčková *et al*. 2017a). Briefly, birds were captured into mist nets when the nestlings were 7-14 days old. Immediately after the capture, they were placed into fresh paper bags for ca. 15–20 min, after which faecal samples were collected using sterile microbial swabs (minitip FLOQSwabs, Copan, Italy) and stored in RNA later at −80 °C. We also measured their tarsus length, weight (for latter calculation of standardized body mass as the ratio of tarsus length and weight) and collected blood and feather samples (not included in this study). The age of the birds was assessed based on differences in plumage coloration of primary and secondary covers (Svensson and Baker, 1992) and bird ringing records. Finally, we ringed all birds with an aluminium ring with an unique code of the Czech Bird Ringing Centre, National Museum in Prague. The research was carried out under the current laws of the Czech Republic and the European Union. The experiment was approved by the Environmental Protection Department of the Prague City Hall (permit No. S-MHMP-1061728/2010/OPP-V-790/R-235/Bu) and the Ministry of the Environment of the Czech Republic (permit No. 22003/ENV/16-1009/630/16). Permission for catching and ringing adult birds was granted by the Bird Ringing Centre of the National Museum in Prague.

### DNA isolation

To maximise our microbial DNA extraction efficiency with minimum contamination risks, we first performed a comparison of five bacterial DNA extraction kits (for details see Supplementary Material and Method online 1, SMMO 1, in Supplementary information 1, SI1). Finally, all microbial DNA samples were extracted using the only DNeasy PowerSoil Kit (Qiagen, Cat. No. 47016, Hilden, Germany) with some modifications (see SMMO 2, SI1) in a laminar biosafety cabinet. The samples were homogenized using vortex with horizontal adapter (cat. no. 13000-V1-24; MO BIO Laboratories, Inc, Carlsbad, USA) for 10 min at maximum spead to optimize DNA isolation efficiency, and the extracted DNA was eluted to 55 μL with elution buffer. As a starting material, for E0-egg 200 μL homogenized egg content was mixed with 200 μL sterile water (to prevent pellet formation), for E13 embryo, ca half of the intestine, and for adult females, whole faecal samples were used. To prevent cross-contamination between different biological sample types, each sample type was extracted separately, strictly abiding by the principles of clean working (decontamination procedures and manipulation minimising the risks of inter-sample contamination). We also included isolation negative controls (INCs) with nuclease-free water (the same batch as for egg content dilution) that were processed separately for each sample type in the following counts: for egg content samples (N = 7), embryonic samples (N = 16), and female faecal samples (N =3).

### Microbial metabarcoding

Our metabarcoding approach was based on amplification of the V3–V4 region of the 16S rRNA gene. For both egg content and embryonic GIT samples, two different protocols were used to maximise the probability of microbiota detection, minimising the effects of primers and amplification kit selection. In protocol 1 (P1), no-chloroplast amplifying primers 335F (CADACTCCTACGGGAGGC) and 769R (ATCCTGTTTGMTMCCCVCRC) (Dorn-In *et al*. 2015) were used together with KAPA2G Robust PCR Kits (Kapa Biosystems, Cat. No. KK5005, Wilmington, USA). In protocol 2 (P2), we used universal bacterial primers S-D-Bact-0341-b-S17 (CCTACGGGNGGCWGCAG) and S-D-Bact-0785-a-A-21 (GACTACHVGGGTATCTAATCC) (Klindworth *et al*. 2013) together with KAPA HIFI Hot Start Ready Mix (Kapa Biosystems, Cat. No. 07958935001) containing the polymerase with proofreading activity. Due to higher proportion of diet-derived chloroplast sequences in female faecal samples, these samples were run only with the no-chloroplast amplifying primers (Kropáčková *et al*. 2017a). Both primer sets were tagged with 10 bp oligonucleotides for multiplexing Nextera™ DNA Sample Prep Kit (Illumina®-Compatible, Cat. No. GA09115, San Diego, USA). For each sample and primer set, PCR was prepared in technical duplicates to check for the consistency of the microbial profiles (see Table S1, SI1 for details on the PCR conditions). Different biological sample types were amplified in different plates to prevent cross-contamination. In each plate, two negative PCR controls (further termed as no PCR template controls, NTCs; RNA free water; Cat. No. 760011596, Qiagen, Hilden, Germany) were used. Then all PCR products were run on 1.5% agarose gel and the PCR product concentrations were assessed based on gel band intensity using GENOSOFT software (VWR International, Belgium). By their concentrations, samples were pooled into several concentration pools and were purified by paramagnetic beads SPRIselect (Beckman Coulter Life Sciences, USA). To remove PCR non-specificities, PCR products in the range of 520-720 bp were excised by Pipin Prep instrument using 1.5% Agarose Cassettes, dye-free, int. Standards (Pippin Prep, 250bp – 1.5kb, Cat. No. 341CDF1503, Biozym, Hessisch Oldendorf, Germany). Subsequently, the concentration of purified pools was checked by Qubit Fluorometer (ThermoFisher Scientific) with Qubit dsDNA BR Assay Kit (ThermoFisher Scientific, Cat. No. Q32850) and pooled at equimolar concentrations. Finally, the resulting amplicon libraries were sequenced using MiSeq Illumina platform with 2L×L300Lbp paired-end reads and v3 chemistry (Illumina) at the Central European Institute of Technology (CEITEC, Brno, Czech Republic). E0-egg and embryonic GIT samples (i.e. the low bacterial biomass samples) were sequenced in different sequencing runs than the female faecal samples (i.e. the high bacterial biomass samples) because DNA concentration levels importantly differed between these biological sample types and feacal samples could cross-contaminate other samples (see Tables S1, SI2 wherein also see for the metadata).

### Bioinformatic and statistical analysis of sequencing data and identification of contaminating taxa

Out of 855 total samples sequenced (number indicated in duplicates), we obtained sequences for all samples with a total of 1 996 026 reads. Samples were then demultiplexed and primers were trimmed by skewer software (Jiang *et al*. 2014). Using dada2 (Callahan *et al*. 2016), we filtered out low-quality sequences (expected number of errors per read less than 2), denoised the quality-filtered fastq files and constructed an abundance matrix representing read counts for individual amplicon sequence variants (ASVs) in each sample. Using uchime (Edgar *et al*. 2011) and gold.fna database (available at https://drive5.com/uchime/gold.fa), we identified chimeric sequences and removed them from the abundance matrix. We aligned the ASV sequences obtained using the two different primer sets (P1 and P2; Table S1, SI1) amplifying slightly different 16S rRNA gene ranges using Decipher R package (Wright 2015) and retained only the region overlapping between the amplicons. The mean amplicon sequence length before trimming was 413 bp (median = 416 bp, min = 389, max = 439) for P1 and 425 bp (median = 427 bp, min = 352, max = 450) for P2, and 404 bp (median = 411 bp, min = 352 bp, max = 416 bp) after trimming. The same sequences after this trimming step were considered as one ASV. Taxonomic assignation of ASVs was conducted by the RDP classifier (80 % confidence threshold, Wang et al., 2007) and Silva reference database (v. 138; Quast et al., 2013). From all downstream analyses, we excluded all ASVs assigned as “Chloroplast”, “Mitochondria”, “Eukaryota” or those that were not assigned to any bacterial phylum. We also removed all samples with a low number of sequences (< 50 sequences with sequencing artefacts or low number of reads not sufficient for statistical analysis). This step reduced 9.59 % of samples and only 0.01 % of all sequence reads (please, see Table S2A-E, SI1 for numbers of sequences/ samples/ ASVs after applying different bioinformatic filtering steps).

For the different sample types and protocols, we assessed the consistency of microbial profiles between technical duplicates using Procrustean analysis based on Bray-Curtis dissimilarities calculated based on ASVs proportions in each sample (Kreisinger *et al*. 2017). Depending on the protocol used, E0 eggs, INC or NTC had lower consistency of technical PCR duplicates than female and embryonic samples (Table S6, SI1). We then removed all ASVs that were not consistently present in both technical duplicates for a given sample (putative contamination and sequencing artefacts). From samples without duplicates, we excluded all ASVs that were presented in just one sample and are not presented in any other duplicated samples. This dramatically reduced both numbers of ASVs (to only 152 out of 1370 ASVs) but only weakly affected the numbers of reads used for the analysis (decline by 9.81 %), suggesting the presence of many lowly abundant ASVs in the non-filtered dataset. Subsequently, read counts for duplicated samples were merged for all later analyses. After combining the sequencing duplicates, we retained 416 samples (out of the total of 773 sequencing reactions).

Importantly, we used the Decontam package (Davis *et al*. 2018) to identify and subsequently eliminate putative contaminating ASVs whose prevalence was elevated in INC and NTC samples compared to all great tit samples and/or that were more prevalent in samples with a low concentration of metagenomic DNA (as assessed by PCR product concentration). The analysis was carried out separately for egg and embryonic samples and for female faecal samples. In total, we removed 12 potentially contaminating ASVs in embryos and egg content samples and 2 in female faeces samples (see Figure S1 and Table S7, SI1 for their full taxonomy) representing 4.84 % drop in sequence reads (please, see Figure 1 and Table S2, SI1 for all filtering steps).

Especially in E0-egg, E13-nat, and E13-PBS samples as well as in the INC samples (i.e. low bacterial biomass samples), the taxonomic composition was dominated by the genus *Ralstonia* which comprised in total of 30.1 % of the sequenced reads. It was also the only one ASVs distributed across all sample types (Figure S7A, SI1 and Table S3A, SI2). This finding together and the fact that *Ralstonia* is known as a frequent contaminant of plastics and solutions (Ryan and Adley 2014) suggest contamination of our samples with this bacterium although it was not detected in Decontam analysis. Despite all of our efforts to keep our procedure clean, *E. faecium* used for *in ovo* treatment might cross-contaminate some other embryonic samples (particularly E13-PBS; Figure S3, SI1). In all subsequent analyses we conservatively present the results with and without *Ralstonia* and *Enteroccocus* included where necessary (please, see the read counts and other parameters for different sample types, Table S2A-E, SI1 before and after their exclusions). After these steps, the final dataset consisted of 191 samples containing sequences, 128 ASVs and 401 089 reads. Finally, we compared these filtered metabarcoding results with the lists of the most commonly contaminating ASVs compiled by Salter *et al*. (2014); Eisenhofer *et al*. (2019) and Stinson, Keelan and Payne (2019).

Using the microeco package, we created Venn diagrams (Liu *et al*. 2021) showing overlaps of ASVs and genera between different biological sample types and PCR protocols. To test whether the female faecal microbiota is more similar to the microbiota of the own egg or embryo than expected by chance, we compared the differences in Jaccard and Bray-Curtis dissimilarities between the female faecal microbiota and the microbiota of the own vs. foreign egg (E0-egg)/embryo (E13-nat) using the Wilcoxon test and P1 dataset. This analysis was carried out for the dataset with *Ralstonia* (N_E0-egg_ = 47, N_E13-nat_ = 66, N_F_ = 30), as there were not enough samples available for statistical analysis after its exclusion. We also tested using the Wilcoxon test if embryos from the same nest are more similar to each other than embryos from different nests in microbial composition. Jaccard dissimilarities were calculated after rarefaction of the abundance matrix (N = 51 sequences per sample, i.e. the minimum number of reads in the dataset). Bray-Curtis dissimilarities were calculated based on relative ASV abundances. All statistical analyses were performed using R software, v.4.1.1 (R Core Team, 2017).

### Quantitative PCR screening of potentially pathogenic bacteria

Based on the metabarcoding results, we developed ASV-specific probe-based 16S rRNA gene (DNA) qPCR assays to detect potentially pathogenic bacteria present in egg content and embryos. We set the following criteria for selection of the target ASVs: (1) the ASV presence in E0-egg as well as in E13-nat samples, (2) the ASV absence from any INCs and NTCs, (3) the ASV being not previously reported as a contaminant of laboratory plastics or chemicals (based on a literature survey), and finally

(4) the ASV being known as a potential pathogen of the gastrointestinal or urogenital tract in animals (pathogens have a greater influence on host physiology and fitness). In cases where the selected ASVs resembled in their sequences other ASVs occurring in INCs or NTCs, we designed the qPCR primers and probes to target only a more dissimilar ASV variant that cannot be non-specifically co-amplified together with any potential contaminant. Passing these criteria, we developed three qPCR assays: i) for *Corynebacterium* (Barbosa and Palacios 2009; Risely *et al*. 2018), ii) for *Clostridium* (Tsiodras *et al*. 2008; Benskin *et al*. 2009) and iii) for *Dietzia* (Koerner, Goodfellow and Jones 2009; Olowookere *et al*. 2022), see Table S3, SI1 for primer and probe sequences.

To increase the sensitivity of qPCR assays, all samples were pre-amplified for 30 cycles with bacterial universal primers (Klindworth *et al*. 2013; the same as in P1) but with high fidelity and accurate polymerase Platinum SuperFi PCR I Master Mix (ThermoFisher Scientific, Cat. No. 12351), see Table S5, SI1 for more details. To minimise DNA binding to the plastic surface, we used low-binding plastics and added 1 µl of 0.1 ng/µL tRNA carrier (i.e. spike water, Qiagen, Cat. No. 1068337) in each tube with stock eluted DNA. Pre-amplification was done in technical triplicates and different biological sample types were run on different plates to minimise the risk of cross-contamination. Three negative pre-amplification controls (NTC-pre-amp; i.e. spike water) were present on each plate. Pre-amplified PCR products were then diluted with spike water 3x and were used as a template for ASV-specific qPCR (see Table S5, SI1 for PCR conditions).

ASV-specific qPCRs were performed with Luna Universal Probe qPCR Master Mix (New England Biolabs, Inc., Cat. No. E3006, Ipswich, Massachusetts, USA) with cycling conditions following manufacturer’s instructions (Table S5, SI1) using the Light Cycler 480 (Roche Applied Science) in a 384-well plate format (Roche Applied Science, Cat. No. 04729749001). All assays were performed in technical triplicates where pre-amplified DNA from technical triplicate was used as a template for independent monoplicate qPCR (allowed by high repeatability between technical triplicates in qPCR). DNA sequences standards (IDT, gBlocks Gene Fragments; Table S4, SI1) in serial dilutions of 10^9^ – 10^1^ were used to estimate the qPCR efficiency (E = 1.995 for *Clostridium*, E = 1.989 for *Corynebacterium* and E = 1.835 for *Dietzia*). Additionally, each plate also contained three template-free negative controls (spike water; NTC-qPCR) and three NTC-pre-amp (as mentioned above).

For each triplicate, we calculated mean C_p_ values using the second derivative method (2^nd^ derivative Max) implemented in LightCycler 480 SW 1.5 (Roche Applied Science). Replicate was considered as positive when C_p_ <36, which corresponds to ∼ 1 – 10 DNA molecules after the pre-amplification (assessed from the gBlock standard curves). However, given the expected high stochasticity of PCR amplification in very low template concentrations, all E0-egg and embryonic GIT samples wherein at least one of the triplicates was positive (C_p_ values <36) were re-assessed with another independent pre-amplification and qPCR (in total, six technical replicates per sample were obtained). Finally, we defined samples as positive only if the C_p_ values were <36 in at least 2/6 replicates and simultaneously, the positivity was confirmed in both independent qPCR runs (see Figure 2 for the procedure overview). Due to the pre-amplification step, we could interpret qPCR results only as semiquantitative, identifying the number of replicates in samples reaching the quantity of bacterial DNA over approximately 1 – 10 molecules per reaction (estimated based on gBlocks standards). Original measured qPCR data are in Table S2, SI1.

Using the qPCR data, we statistically tested whether *Corynebacterium, Clostridium* and *Dietzia* occur in biological samples more frequently than in negative controls (INC, NTC-qpCR and NTC-pre-amp). We applied Generalized Linear Models (GLMs) with quasibinomial distribution where the ratio of the number of positive replicates (C_p_ < 36) to the total number of replicates was a dependent variable and biological sample type was an independent variable. Separate models were built for each combination of the assay (i.e. given ASV) and biological sample type (here E0-nat, E13-nat and F) except the cases where given ASV was not confirmed by the independent qPCR in any sample (e.g. for *Clostridium* or *Corynebacterium* in E0-nat and E13-nat; please, see all models M1-5 in SI3). Plots were generated using ggplot2 (Wickham 2016), boot (Canty and Ripley 2021) and ggeffects packages (Lüdecke 2018). All statistical analyses were performed using R software, v.4.1.1 (R Core Team, 2017).

## Results

Our results on microbiota composition obtained using the two metabarcoding protocols were generally highly consistent. Since the P1 data contain fewer sequencing artefacts, we primarily show the results of the P1 approach here and provide the P2 results as well as data with *Ralstonia* and *Enterococcus* in SI for comparison. However, in cases where a comparison of data is required or where there are discrepancies between the P1 and P2 results, we highlight this also in the main text.

### (i) Microbial profiling in egg content, embryos and females

Applying various bioinformatic steps greatly reduced number of samples, sequences as well as ASV diversity and further parameters in low bacterial biomass samples (see Figure 3 and S2A-E, SI1 for sample-specific statistics). For instance, out of initial 1 996 026 reads, the pre-final dataset had 1 711 376 reads but the only 401 089 reads remained in the final dataset when *Ralstonia* and *Enterococcus* were excluded, of which 1236 reads (0.06 % of initial reads) belonged to E0-egg, 45 488 reads (2.28 %) to E13-nat and 272 337 reads (13.64 %) belonged to female faecal samples. This indicates that despite the increased number of PCR cycles in E0-egg and E13-nat samples, both E0-egg and E13-nat samples contained only very few bacterial sequences and such microbial data are highly atypical. In eggs (E0-egg), when we excluded *Ralstonia* out of 52 samples only 8 samples had sufficient number of sequences (mean = 154.5 reads per sample) and 84.6 % of samples did not contain any bacterial sequences after the filtering (data obtained using P1; Table S2A-E and Figure S2, SI1). In total, only 11 ASVs were found with mean 2.38 ASV per sample (Table S2D and E, SI1). Specifically, among the most common ASVs in the E0-egg samples family of Xanthobacteraceae and genera of *Cutibacterium* and *Mycobacterium* were detected in at least five samples and further *Sphingomonas* was detected in two samples (Table 1; Figure 4; Figure S3 and Figure S4, SI1). *Clostridium* was revealed in one E0-egg sample. P2 revealed the similar ASV numbers as P1 (15) but included two putative pathogens, *Dietzia* (one sample) and *Corynebacterium* (three samples).

**Figure 3:**
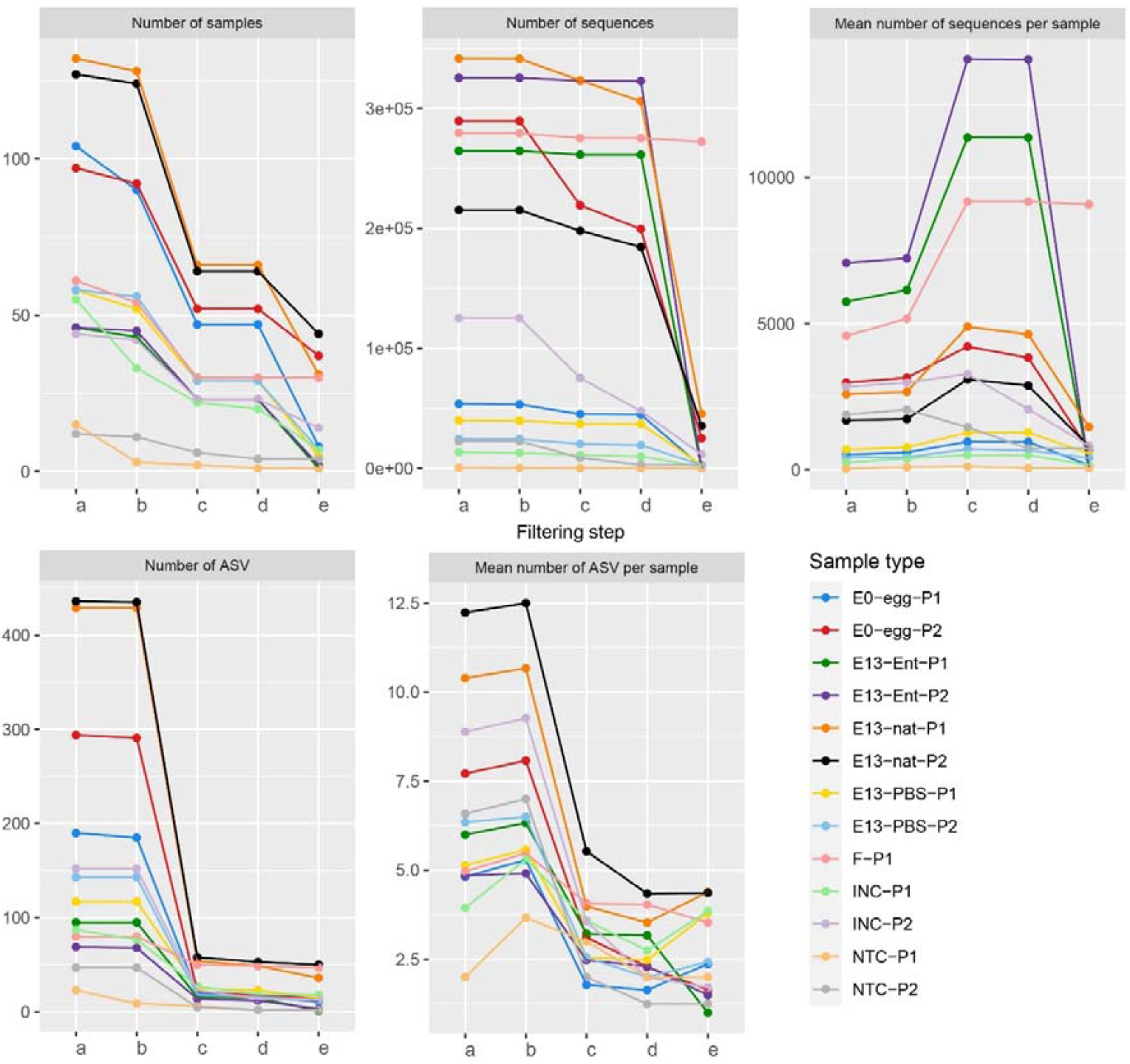
Changes in the numbers of samples, sequences, and bacterial amplicon sequence variants (ASVs) after applying different bioinformatic filtering steps. Different bioinformatic steps were applied: a – all samples after sequencing, b – removing samples with a low number of sequences (< 50; the same limit applied also to steps c-e), c – removing samples with inconsistent technical duplicates, d – removing contaminants by Decontam, e – removing *Ralstonia*/ *Enterococcus*. The comparison is shown for different amplification protocols (P1 and P2, see methods for more details). Please note that the characteristics for steps a-c are given in technical duplicates (i.e. real number of samples * 2). E0-egg – egg content sample at embryonic day 0, E13-nat – E13 intestinal sample from non-manipulated egg, E13-Ent – E13 intestinal sample from *Enterococcus*-treated egg, E13-PBS – E13 intestinal sample from a control PBS-injected egg, F – adult female faecal sample (amplified only in P1), INC – negative control of isolation (i.e. isolation negative control), NTC – negative control of PCR (no template PCR control). For specific values in all these steps, please, see Tables S2A-E, SI1.

**Figure 4:**
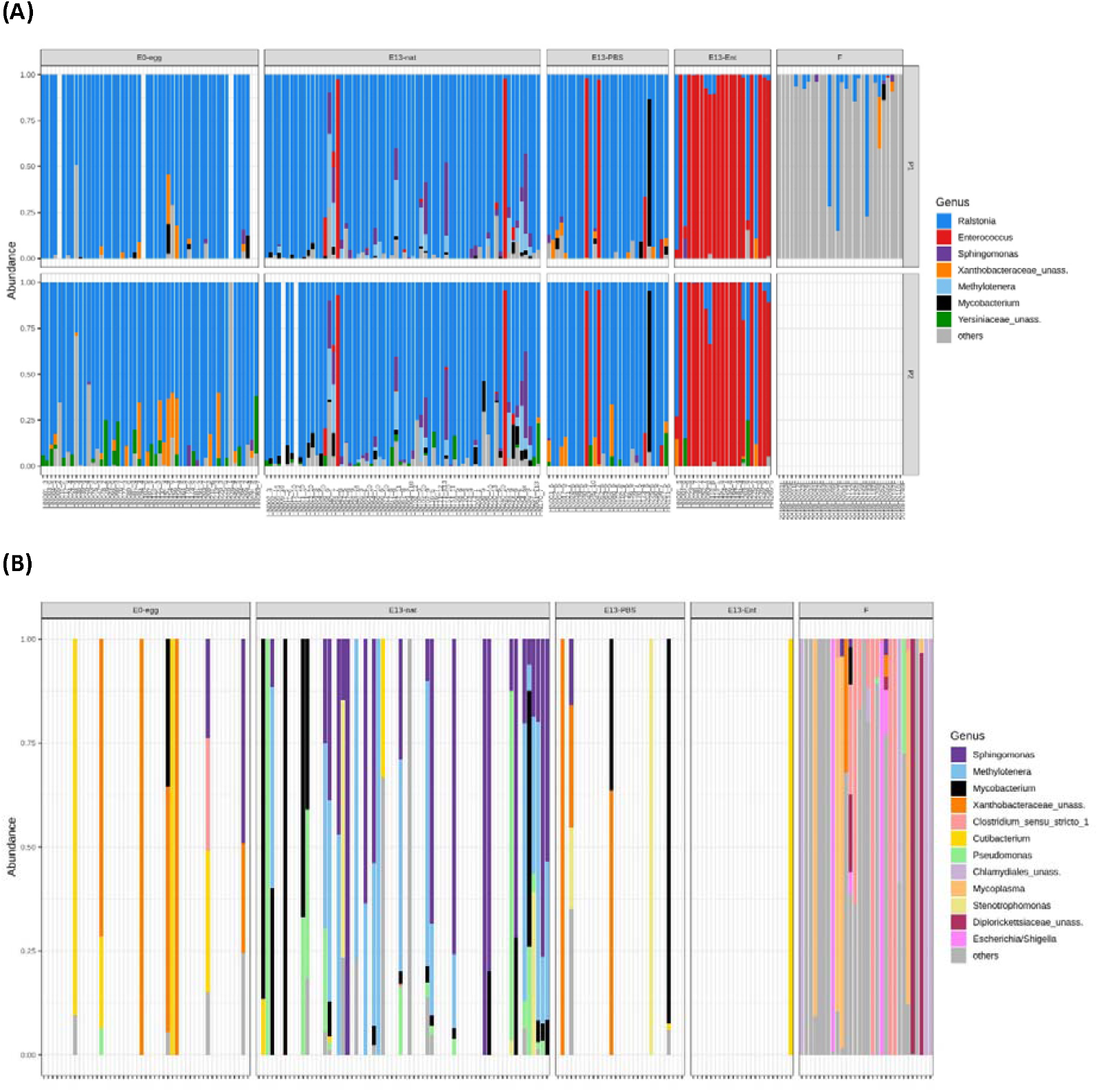
Relative abundances of bacterial taxa (A) before excluding potentially contaminating *Ralstonia* and cross-contaminating *Enterococcus* sequences for both protocols (P1 and P2) and (B) and after their exclusion in great tit egg and embryonic samples only for P1. Only the most abundant genera (relative abundance > 1 %) are shown. Less abundant genera are included in the category ‘Others’. Results for different amplification protocols (P1 and P2, see methods for more details) are shown. E0-egg – egg content sample at embryonic day 0, E13-nat – E13 intestinal sample from non-manipulated egg, E13-Ent – E13 intestinal sample from a *Enterococcus*-treated egg, E13-PBS – E13 intestinal sample from a control PBS-injected egg, F – female faecal samples (amplified only by P1). For the relative abundances in P2 after Ralstonia and *Enterococcus* exclusion, see Figure S4, S1.

**Table 1:**
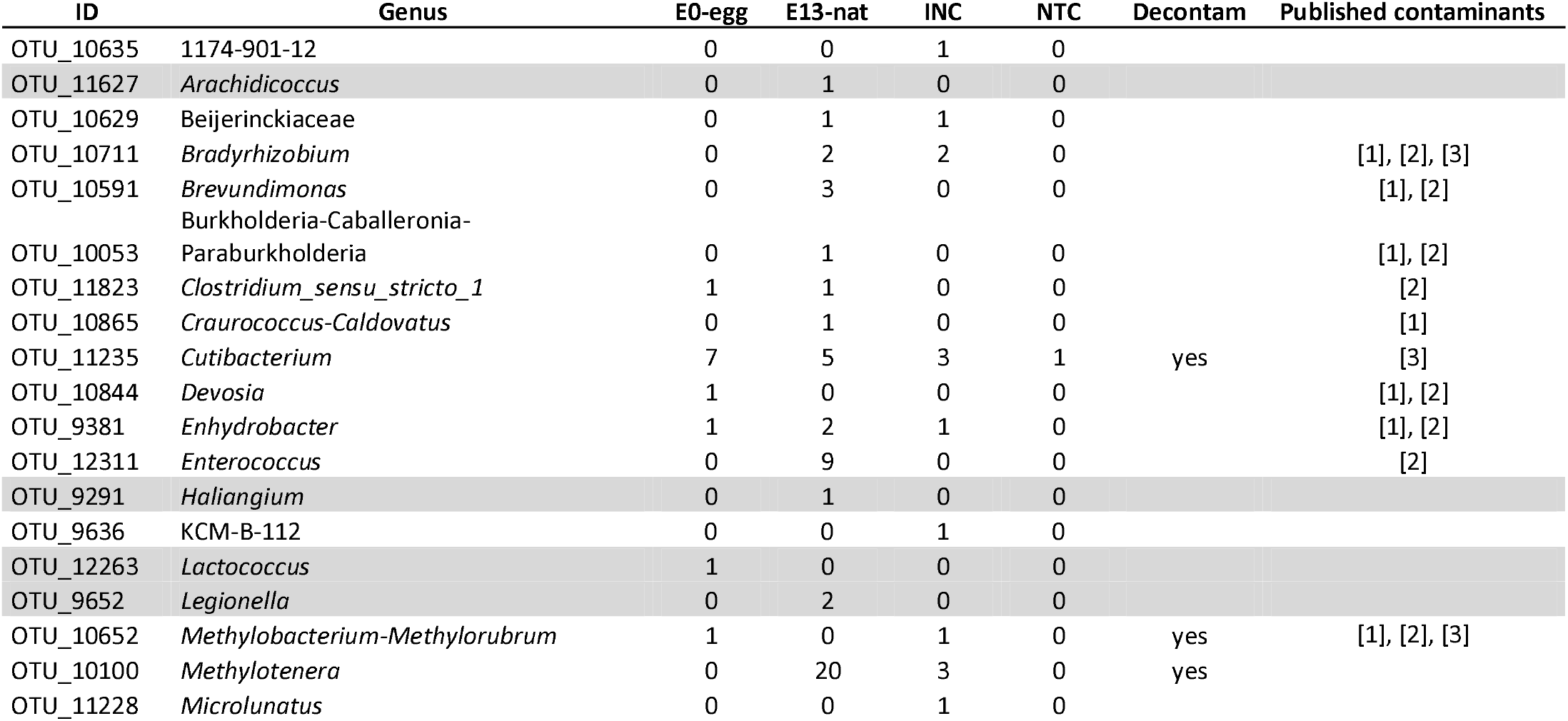
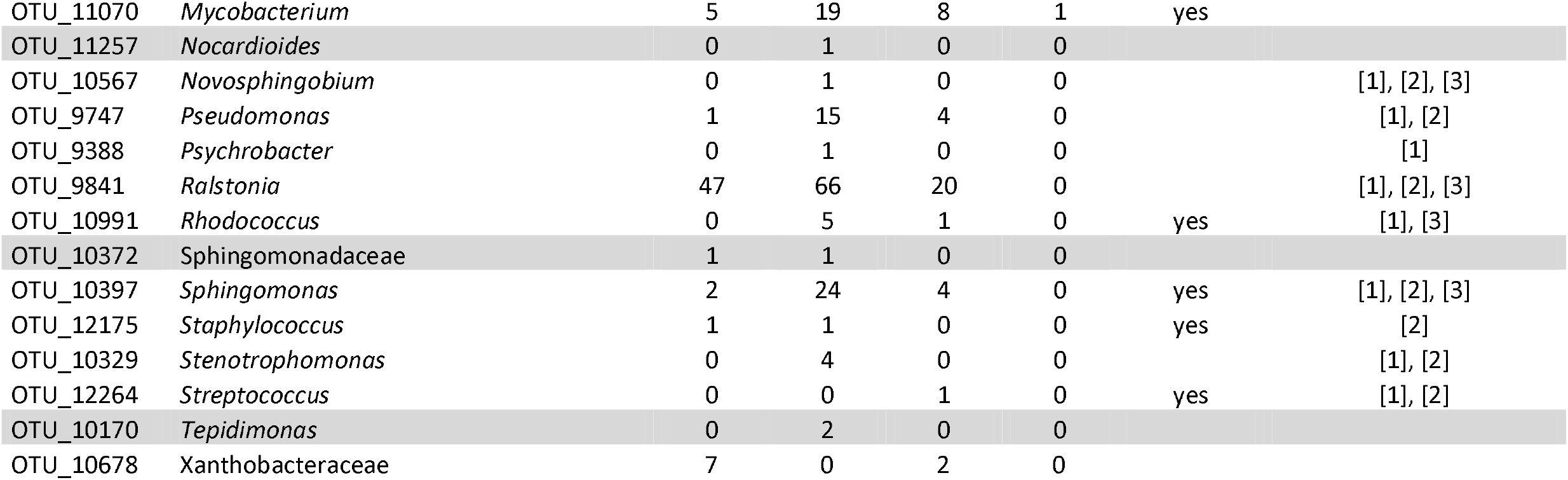
List of bacterial genera found in egg content (E0-egg) and embryonic intestine samples (E13-nat) in the great tit. The lowest taxonomic assignments into genera, where possible, are shown based on the RDP classifier (Wang et al., 2007) and Silva reference database (Quast et al., 2013). For a complete list of all amplicon sequence variant (ASVs) and genera in all sample types, please see Table S3A and B, SI2). E0-egg – egg content samples at embryonic day 0 (N = 47), E13-nat – embryonic intestine sample at embryonic day 13 (N = 66), INC – negative controls of isolation (N = 20), NTC – negative control of PCR (N = 1). Number in brackets indicates the number of samples in given category after bioinformatic filtering before removing *Ralstonia* and *Enterococus*. Presence of given taxa for protocol 1 (P1; see methods for more details) is indicated by the number of positive samples. Decontam – genera revealed in the statistical analysis by Decontam are indicated (for a complete list of all contaminating ASVs revealed by Decontam analysis, please see Table S7, SI1). Previously reported genera as common contaminants are cited: [1] – Salter *et al*. 2014, [2] – Eisenhofer *et al*. 2019, [3] – Stinson, Keelan and Payne 2019. Genera that were only identified in E0-egg and E13-nat samples and not in negative controls are highlighted in grey.

In embryonic GIT (E13-nat) out of 66 samples only 31 had enough sequences after all filtering steps (mean = 1467.4 reads per sample) and 53.03 % samples did not contain any sequences (Table S2A-E and Figure S2, SI1). In embryonic E13-nat samples, we revealed higher ASV diversity than in E0-eggs, with a total of 49 ASVs and mean 4.39 ASV per sample (Table S2D and E, SI1). Out of these, 28 ASVs were unique to E13-nat, covering, however, only 2.3 % of all reads (Figure S7A and see further panels B-E for comparisons on genus-level and also for P2, SI1). The E13-nat samples were dominated by genera *Sphingomonas, Mycobacterium, Methylotener* and *Pseudomonas* (found in > 10 samples) and further by *Cutibacterium, Stenotrophomonas, Enhydrobacter, Legionella*, Blastocatellia, *Brevundimonas* and *Tepidiomonas* (found in ≥ 2 samples; Table 1; Figure 4; Figure S3 and Figure S4, SI1). *Clostridium* was revealed in one E13-nat sample. Additionally, using the P2 approach, we again revealed the two putative pathogens, *Dietzia* (two samples) and *Corynebacterium* (one sample). Out of the total 49 ASVs detected in the E13-nat samples, 9 ASVs were shared with the E0 eggs, including the most common genera, such as *Cutibacterium, Mycobacterium, Spingomonas* (Table 1 and Figure S7A and also B-E for further comparisons, SI1).

The embryonic GIT samples originating from E0 eggs experimentally injected with *E. faecium* (E13-Ent) were, as predicted, dominated by the genus *Enterococcus*, independent of the dosage applied (Figure S3 and S5, SI1). This demonstrates that our 16S rRNA gene metabarcoding approach is sensitive to the detection of bacteria even when they are present in low quantities in the eggs (≥ 1000 copies, i.e. the low dose of *E. faecium. In ovo* treatment also did not affect the microbiome of E13, as the control E13-PBS samples contained a similar composition of ASVs as E13-nat samples (Figure 3 and Table S8, SI1). Despite all our efforts to keep our procedure contamination-free, we observed a high abundance of *Enterococcus* in a few E13-PBS samples, possibly due to cross-contamination.

Female faecal samples had generally a much higher number of reads per sample than the E0-egg and E13-nat samples (Table S2A-E, SI1). In total, 30 female faecal samples contained 49 ASVs, out of which 34 ASVs were unique only to the faeces (this covered 28.7 % of all sequences, Figure S7A, SI1). We observed high inter-individual variability between females (Figure S6A and B, SI1) and taxonomically different composition from all other biological sample types (Figure 4). Among the most common taxa, *Mycoplasma, Clostridium, Escherichia, Lactococcus*, Diplorickettsiacae or *Ureoplasma* were detected. There were only six ASVs shared between E0-egg and female samples (e.g. in genera *Clostridium, Devosia gracialis, Lactococus, Mycobacterium Sphingomonas*) and 9 ASVs shared between E13-nat and female samples (e.g. in genera *Mycobacterium, Rhodococcus*, Sphingomonas; Table S3A,B, SI2 and see Figure S7A, SI1).

### (iii) Similarity in microbial communities within- and between nests

Female microbiota was not more similar to the microbiota of their own eggs (E0-egg) compared to foreign eggs (for Jaccard W = 15174, *p* = 0.726 and Bray-Curtis W = 15152, *p* = 0.741). On the other hand, we detected slightly increased similarity between female faecal microbiota and microbiota of embryos in their nests (E13-nat) than would be expected by chance, which was the case for Jaccard (W= 39346, *p* = 0.086), but not for Bray-Curtis (W= 36823, *p* = 0.368). E13-nat within the same nest were not more similar in microbial composition than E13-nat between different nests (for Jaccard W = 21862, *p* = 0.021 and Bray-Curtis W= 35762, *p* = 0.125).

### (iv) qPCR detection of potentially pathogenic bacteria

Unlike in 16S rRNA gene metabarcoding, no *Corynebacterium* and *Clostridium* have been found by specific qPCR assays in any of the 52 E0-eggs samples (see Table S8, SI1 for comparison of NGS and qPCR results). In contrast, *Dietzia* was revealed by the qPCR in four E0-egg samples (7.62 %). However, *Dietzia* was detected also in INC and its frequency in E0-eggs statistically did not differ from any type of the negative controls in E0-egg dataset (Sample type: *p* = 0.318, Model 1, M1 in SI3 wherein see for full model details, and Figure S8A, SI1).

Also, in E13-nat samples, we detected no *Corynebacterium* and *Clostridium* in any of the 66 E13-nat samples assessed. *Dietzia* was found by the qPCR only in two E13-nat samples (i.e. 2.66 %). Overall *Diezia* positivity did not statistically differ from negative controls (especially from INC) in the E13-nat dataset (Sample type: *p* = 0.154, M2, SI3 and Figure S8B, SI1).

In female faeces, all *Corynebacterium* (25.00 % of samples), *Clostridium* (90.63 %) and *Dietzia* (90.63 %) have been detected. However, when further statistically tested, only *Clostridium* and *Corynebacterium* had significantly higher positivity in the faecal samples compared to the negative controls (Sample types: *p* = 0.004, M3 and *p* = 0.042, M4, SI3 and Figure S8C and D, respectively) and not *Dietzia* (for Sample type: *p* = 0.103, M5, SI3 and Figure S8E, SI1). Unlike in low bacterial biomass samples (i.e. E0-egg and E13-nat), virtually all *Clostridium*-positive samples (i.e. 91.66 %) revealed by NGS in females were also confirmed by the more sensitive qPCR, indicating that for the higher bacterial biomass samples both methods give consistent results (Table S8, SI1).

## Discussion

Using combinations of two 16S rRNA-based metabarcoding protocols and specific qPCR assays, we described in this study microbiome of the egg content, embryonic intestine and mother female faeces in a free-living passerine, the great tit. Importantly, unlike in previous avian studies, we performed our PCR in technical replicates and prepared the sequencing runs for low and high bacterial biomass samples independently to minimise the risks of amplification bias and possible cross-contamination. Contrary to some previous studies in birds (e.g. Ding et al., 2017; Trevelline et al., 2018), our results revealed negligible and non-consistent microbiota communities in eggs, with a high proportion of potential contaminants. In embryonic samples, there were more bacterial ASVs, yet still these frequently represented putative contaminants. Out of three potentially pathogenic ASVs detected by our NGS metabarcoding in the E0-egg and E13-nat samples (*Corynebacterium, Clostridium* and *Dietzia*), only *Dietzia* was confirmed by qPCR in E0-egg and E13-nat, which might support the putative *vertical transfer*.

E0-egg samples had only negligible traces of microbiota as documented by the high proportion of samples without any sequences after the filtering and low mean ASV and number of reads per sample despite the increased number of PCR cycles. The egg-content microbial communities were dominated by Xanthobacteraceae, *Cutibacterium, Mycobacterium* and *Sphingomonas*. However, virtually all ASVs identified in E0-eggs were also detected in our INCs and/ or NTCs and most of them are also known as frequent contaminants of laboratory materials (see discussion below and Table 1 for the detailed list of contaminants). Also, our results indicate that the E0 microbiota in eggs from the same nest was not more similar than the one between eggs from different nests. Unlike for *Dietzia*, no *Corynebacterium* and *Clostridium* were detected by qPCR in any E0-egg samples, which supports our nearly sterile egg hypothesis in great tit. This also highlights the importance of validating sequencing results with the more accurate qPCR. Our results suggest only negligible microbiota in great tit eggs at the E0 and are, thus, mostly consistent with the *Sterile egg hypothesis*. This is supported by (i) the absence of clearly distinguishable microbiota between the E0-egg and NTC and INC samples, (ii) non-consistency of technical duplicates, (iii) low ASV abundance and diversity and (iv) the high proportion of potential contaminants. The only exception supporting the possibility of vertical transfer in our data might represent *Dietzia*, which was consistently identified in female faeces and in egg and embryonic samples (yet occurred also in negative controls). Contrary to our results, in E0 egg whites of the Korean chickens, diversified bacterial communities were previously revealed, with the dominance of *Pseudomonas* (65 % of all reads), *Janthinobacterium*, Burkholderiales, *Flavobacterium, Stenotrophomonas, Acinetobacter*, Enterobacteriaceae, Comamonadaceae and Xanthomonadaceae (Lee *et al*. 2019). Microbial communities in egg whites (E0) distinct from the negative controls were found also in four passerine species, yet in a very small dataset of only 11 eggs in total (Trevelline *et al*. 2018). Different egg microbial communities reported at the E0 stage between these studies might either indicate high inter-species variability or reflect insufficient control for the stochasticity of sequencing data (as they did not perform PCR in technical duplicates).

The risk of potential contamination is a critical issue in low-bacterial biomass sample studies (Eisenhofer *et al*. 2019; Salter *et al*. 2014). The sample contamination may originate from different routes, such as the environment where samples are collected, from DNA extraction kits, PCR chemicals and plastics, wet lab procedures or cross-contamination between different samples (Eisenhofer *et al*. 2019). Unlike in mammalian studies where an ongoing debate whether human placenta and foetus are sterile (Perez-Muñoz *et al*. 2017) or not (Aagaard *et al*. 2014) has stimulated the usage of very careful protocols to assess the contamination risk (de Goffau *et al*. 2019), in birds, unfortunately, these caveats have been only rarely addressed. For instance, in chicken egg white and embryonic microbiota studies, no negative controls were integrated (Ding *et al*. 2017, 2022) or these were of only insufficient types and numbers (Akinyemi *et al*. 2020) that limits their inferences of microbiota profiles (but see Grond et al., (2017) and Lee et al., (2019). Not surprisingly, the most abundant taxon in the chicken embryonic samples, *Halomonas* (Ding *et al*. 2017), was later indicated as a saline buffer contaminant of the DNA isolation kit (Lee *et al*. 2019). Our study is the first describing bacterial recruitment in bird embryos by adopting multiple protocols or the technical duplicate-based approach. Incorporating these measures and various bioinformatic filtering steps helped us to detect PCR amplification bias, which may be significant in low bacterial biomass and to remove many inconsistent ASVs that likely originated from environmental contaminants or as cross-contaminants. While the initial dataset consisted of 1382 ASVs, it was first reduced to 1370 ASVs after removing samples with low number of sequences (< 50), then dramatically reduced to 152 ASVs after removing samples with non-consistent technical duplicates and finally to 128 ASVs after removing statistically identified contaminants from Decontam analysis and *Ralstonia* and *Enterococcus*. Focusing on the final dataset of E0-egg and E13-nat samples in the great tits, we revealed in total 29 genera. In contrast, in studies without these measures even up to one magnitude higher ASV diversity was found in GIT of chicken embryos (Akinyemi *et al*. 2020; Ding *et al*. 2022). Despite that, we still did find some potentially contaminating ASVs based on the literature survey. Contaminating ASVs in previous microbial studies have been mostly soil- or water-dwelling bacteria and bacteria frequently associated with nitrogen fixation (Salter *et al*. 2014). From these known contaminants (Salter *et al*. 2014; Eisenhofer *et al*. 2019; Stinson, Keelan and Payne 2019), we identified e.g. *Pseudomonas, Ralstonia, Rhodococcus, Sphingomonas* and *Stenotrophomonas* in our E0-egg and E13-nat samples. Especially *Ralstonia*, a common contaminant of the DNA isolation kits that can even pass through a 20 nm bacterial filter (Sundaram *et al*. 1999), was highly dominating in our low bacterial biomass samples, comprising in total 30.1 % of the sequence reads. Other contaminants, such as *Propionibacterium* or *Streptococcus* are common human skin-associated organisms (Byrd, Belkaid and Segre 2018). Yet, although the vast majority of the ASVs observed in our E0-egg and E13-nat samples are putative contaminants, we cannot exclude the possibility that they also play some biological roles in the eggs and developing embryos, as some of them have been revealed in bird GIT (e.g. *Mycobacterium* and *Pseudomonas*; Kropáčková *et al*. 2017b, 2017a). Therefore, we can only conclude that their presence in the avian eggs is highly speculative. In addition, some identified genera are also known to be opportunistic pathogens in humans, such as some members of the above-mentioned genera *Mycobacterium* (Primm, Lucero and Falkinham 2004) or *Pseudomonas* (Kerr and Snelling 2009). Unfortunately, with our V3-V4 16S rRNA gene metabarcoding approach, we could not distinguish most ASVs at a lower taxonomic level than genera and the potential pathogenicity of some ASVs could also be highly species-, strain-or condition-specific (Pan, Yang and Zhang 2014).

Compared to E0-egg, the E13-nat microbiome was slightly more consistent, with higher ASV diversity and the number of reads per sample, yet 53.03 % of samples did not obtain any sequences after the data quality filtering. Out of the 49 ASVs detected in the E13-nat samples, 9 ASVs were shared with the E0 eggs, including the most common genera, such as *Cutibacterium, Mycobacterium, Spingomonas*. Yet, again, all genera in E13-nat except for *Stenotrophomonas, Legionella*, Blastocatellia, and *Tepidiomonas* were detected also in our INC and/ or NTC. Also, the absence of higher microbial similarity in E13-nat within the nests than between the nests indicated rather stochastic recruitment of microbiota at E13. From genera found in our great tit E13-nat samples, five genera also inhabited the chicken embryos (E19; Ding et al., 2017): *Pseudomonas, Sphingomonas, Bradyrhizobium, Rhodococcus* and *Staphylococcus*. In another study (Lee *et al*. 2019), four out of eight most abundant taxa in E18 chickens: *Pseudomonas*, Burhkhodriales, *Stenotrophomonas* and Enterobacteriaceae were also present in the great tit E13-nat samples. Despite the sharing of these few genera between the chicken and great tit embryos, overall abundancies and taxonomical diversities in the great tit embryos were much lower than in the chicken embryos (Lee *et al*. 2019; Ding *et al*. 2017, 2022). On the one hand, this could reflect interspecific differences, as the embryos of the altricial great tit are less developed than those of the precocial chickens, on the other hand, the differences in the results may also be due to differences in the NGS protocols used (see above). Our results, therefore, better correspond with those reported by Grond *et al*. (2017) who detected in two arctic superprecocial shorebirds: the dunlin (*Calidris alpina*) and the sandpiper (*Calidris pusilla*) by NGS and qPCR only negligible microbiota in the embryonic GIT just before hatching, with stochastic communities skewed towards dominance of *Clostridia* and *Gammaproteobacteria*.

Using metabarcoding we detected potentially pathogenic genera *Corynebacterium, Clostridium* and *Dietzia* both in E0-egg and E13-nat, and partially also in female faecal samples, suggesting their putative *vertical transfer*. Unlike the sequencing, no *Corynebacterium* and *Clostridium* were detected by qPCR both in E0-egg and E13-nat. Importantly, *Dietzia* was revealed by qPCR in 7.69 % of E0-eggs and in 3.03 % of E13-nat samples. This suggests that *Dietzia* might be in very low frequencies vertically transmitted. Similarly, the putative vertical transmission of pathogenic bacteria from infected mothers to their eggs was suggested both in domestic (e.g. for *Salmonella* in chickens; Keller *et al*. 1995; Gantois *et al*. 2009; Pedroso 2009) and wild birds (e.g. for *Nesseria* in the greater white-fronted geese, *Anser albifrons*; Hansen *et al*. 2015). Since we found almost no microbiota in E0-egg samples but found slightly more diversified microbiota in E13-nat (with limited overlap with E0-egg samples), we think that *bacterial trans-shell migration* in E13-nat seems to have a stronger effect than *vertical transfer*, yet it is still rare in the great tit. As described previously (Kropáčková *et al*. 2017a), we found rich bacterial communities in the female faecal microbiome. However, the fact that these were taxonomically very different from those in the E0-egg and E13-nat samples might suggest that the original embryonic ASVs are either replaced by other ASVs after hatching or exist in the adults only in very low, undetectable amounts. Our results thus suggest that GM is predominantly formed after hatching in passerines. This is also supported by the high ontogenetic dynamics of passerine GM during the nestling period (e.g. Chen et al., 2020; Kreisinger et al., 2017; Teyssier et al., 2018).

## Conclusion

In our study combining 16S sRNA gene metabarcoding with targeted qPCR, we have revealed only nearly sterile egg content and GIT of the developing embryos in the great tit. Our results suggest that out of three ASVs further targeted by qPCR, only *Dietzia* might be passed by *vertical transfer* from mother to egg. We found more support for *bacterial trans-shell migration* during embryogenesis by forming simple and low-abundant bacterial communities in some (but not all) tit embryos. Yet, again, this effect was weak suggesting that GIT microbiota must be dominantly formed in passerines only after hatching. Further careful investigation of egg and embryonic microbiota across avian phylogeny is needed to see whether the described discrepancy in microbiota recruitment between passerines and chickens reflects differences in species life history (e.g. altricial vs precocial nestlings) or rather methodological differences between studies. Our study highlights the importance of using technical PCR duplicates and internal controls to eliminate stochastic noise and contamination in sequencing data which are particularly common in studies with low microbial biomass. Our results indicate that further studies in species with putatively abundant and diversified microbial communities in the embryonic intestine (e.g. chickens) should determine whether bacteria in bird eggs are alive (e.g. by labelling dead/ live bacteria) and metabolically active (e.g. by metatranscriptomics) and whether they have an impact on immune gene expression. Elucidating the mechanisms of GIT microbiota establishment in early ontogeny can improve our understanding of parental effects, contributing both to basic knowledge of wildlife evolutionary ecology as well as to zoohygiene and veterinary applications that minimise risks of disease transmission.

## Acknowledgement

We thank Dan Divín and Jiří Eliáš for their help with field sampling, Dagmar Čížková, Anna Bryjová and Zuzana Świederská for their help and advice with molecular-genetics analysis.

## List of supplementary materials

SI1 – Supplementary methods, figures and tables (main supplement)

SI2 – Metabarcoding and qPCR sample metadata, metabarcoding abundance table for amplicon sequence variants (ASVs) and genera

SI3 – Full Generalized Linear Models (GLMs) details

## References

Aagaard K, Ma J, Antony KM et al. The Placenta Harbors a Unique Microbiome. Sci Transl Med May 2014;21:237–65.

Akinyemi FT, Ding J, Zhou H et al. Dynamic distribution of gut microbiota during embryonic development in chicken. Poult Sci 2020;99:5079–90.

Bäckhed F, Ley RE, Sonnenburg JL et al. Host-bacterial mutualism in the human intestine. Science (80-) 2005;307:1915–20.

Barbosa A, Palacios MJ. Health of Antarctic birds: A review of their parasites, pathogens and diseases. Polar Biol 2009;32:1095–115.

Benskin CMWH, Wilson K, Jones K et al. Bacterial pathogens in wild birds: A review of the frequency and effects of infection. Biol Rev 2009;84:349–73.

Bruce J, Drysdale EM. Trans-shell transmission BT - Microbiology of the Avian Egg. In: Board RG, Fuller R (eds.). Microbiology of the Avian Egg. Boston, MA: Springer US, 1994, 63–91.

Byrd AL, Belkaid Y, Segre JA. The human skin microbiome. Nat Rev Microbiol 2018;16:143–55.

Callahan BJ, McMurdie PJ, Rosen MJ et al. DADA2: High-resolution sample inference from Illumina amplicon data. Nat Methods 2016;13:581–3.

Campos-Cerda F, Bohannan BJM. The Nidobiome: A Framework for Understanding Microbiome Assembly in Neonates. Trends Ecol Evol 2020;35:573–82.

Canty A, Ripley BD. boot: Bootstrap R (S-Plus) Functions. 2021.

Chen CY, Chen CK, Chen YY et al. Maternal gut microbes shape the early-life assembly of gut microbiota in passerine chicks via nests. Microbiome 2020;8:1–13.

Cook MI, Beissinger SR, Toranzos GA et al. Incubation reduces microbial growth on eggshells and the opportunity for trans-shell infection. Ecol Lett 2005;8:532–7.

Cuperus T, Coorens M, van Dijk A et al. Avian host defense peptides. Dev Comp Immunol 2013;41:352–69.

Davis NM, Proctor DiM, Holmes SP et al. Simple statistical identification and removal of contaminant sequences in marker-gene and metagenomics data. Microbiome 2018;6:1–14.

Ding J, Dai R, Yang L et al. Inheritance and establishment of gut microbiota in chickens. Front Microbiol 2017;8:1–11.

Ding P, Liu H, Tong Y et al. Developmental change of yolk microbiota and its role on early colonization of intestinal microbiota in chicken embryo. Animals 2022;12:1–15.

Dorn-In S, Bassitta R, Schwaiger K et al. Specific amplification of bacterial DNA by optimized so-called universal bacterial primers in samples rich of plant DNA. J Microbiol Methods 2015;113:50–6.

Edgar RC, Haas BJ, Clemente JC et al. UCHIME improves sensitivity and speed of chimera detection. Bioinformatics 2011;27:2194–200.

Eisenhofer R, Minich JJ, Marotz C et al. Contamination in Low Microbial Biomass Microbiome Studies: Issues and Recommendations. Trends Microbiol 2019;27:105–17.

Funkhouser LJ, Bordenstein SR. Mom Knows Best: The Universality of Maternal Microbial Transmission. PLoS Biol 2013;11:1–9.

Gantois I, Ducatelle R, Pasmans F et al. Mechanisms of egg contamination by Salmonella Enteritidis: Review article. FEMS Microbiol Rev 2009;33:718–38.

de Goffau MC, Lager S, Sovio U et al. Human placenta has no microbiome but can contain potential pathogens. Nature 2019;572:329–34.

Grond K, Lanctot RB, Jumpponen A et al. Recruitment and establishment of the gut microbiome in arctic shorebirds. FEMS Microbiol Ecol 2017;93:1–9.

Hansen CM, Meixell BW, Van Hemert C et al. Microbial infections are associated with embryo mortality in arctic-nesting geese. Appl Environ Microbiol 2015;81:5583–92.

Honda K, Littman DR. The microbiome in infectious disease and inflammation. Annu Rev Immunol 2012;30:759–95.

Jiang H, Lei R, Ding SW et al. Skewer: A fast and accurate adapter trimmer for next-generation sequencing paired-end reads. BMC Bioinformatics 2014;15:1–12.

Keller LH, Benson CE, Krotec K et al. Salmonella enteritidis colonization of the reproductive tract and forming and freshly laid eggs of chickens. Infect Immun 1995;63:2443–9.

Kerr KG, Snelling AM. Pseudomonas aeruginosa: a formidable and ever-present adversary. J Hosp Infect 2009;73:338–44.

Kizerwetter-Świda M, Binek M. Bacterial microflora of the chicken embryos and newly hatched chicken. J Anim Feed Sci 2008;17:224–32.

Klindworth A, Pruesse E, Schweer T et al. Evaluation of general 16S ribosomal RNA gene PCR primers for classical and next-generation sequencing-based diversity studies. Nucleic Acids Res 2013;41:1–11.

Koerner RJ, Goodfellow M, Jones AL. The genus Dietzia: A new home for some known and emerging opportunist pathogens. FEMS Immunol Med Microbiol 2009;55:296–305.

Kreisinger J, Kropáčková L, Petrželková A et al. Temporal stability and the effect of transgenerational transfer on fecal microbiota structure in a long distance migratory bird. Front Microbiol 2017;8:1–19.

Kropáčková L, Pechmanová H, Vinkler M et al. Variation between the oral and faecal microbiota in a free-living passerine bird, the great tit (Parus major). PLoS One 2017a;12, DOI: 10.1371/journal.pone.0179945.

Kropáčková L, Těšický M, Albrecht T et al. Codiversification of gastrointestinal microbiota and phylogeny in passerines is not explained by ecological divergence. Mol Ecol 2017b;26:5292–304.

Lee SW, La TM, Lee HJ et al. Characterization of microbial communities in the chicken oviduct and the origin of chicken embryo gut microbiota. Sci Rep 2019;9:1–11.

Liu C, Cui Y, Li X et al. Microeco: An R package for data mining in microbial community ecology. FEMS Microbiol Ecol 2021;97:1–9.

Lüdecke D. ggeffects: Create Tidy Data Frames of Marginal Effects for “ggplot” from Model Outputs. 2018.

Lunam CA, Ruiz J. Ultrastructural analysis of the eggshell: Contribution of the individual calcified layers and the cuticle to hatchability and egg viability in broiler breeders. Br Poult Sci 2000;41:584–92.

Mann K. The chicken egg white proteome. Proteomics 2007;7:3558–68.

Olowo-okere A, Ibrahim YKE, Lo CI et al. Bhargavaea massiliensis sp. nov. and Dietzia massiliensis sp. nov., Novel Bacteria Species Isolated from Human Urine Samples in Nigeria. Curr Microbiol 2022;79:1–8.

Ost KS, Round JL. Communication Between the Microbiota and Mammalian Immunity. Annu Rev Microbiol 2018;72:399–422.

Pan X, Yang Y, Zhang JR. Molecular basis of host specificity in human pathogenic bacteria. Emerg Microbes Infect 2014;3:0.

Pedroso AA. Which came firstL: the egg or its microbiotaL? Poult Inf Prof 2009:1–5.

Perez-Muñoz ME, Arrieta MC, Ramer-Tait AE et al. A critical assessment of the “sterile womb” and “in utero colonization” hypotheses: Implications for research on the pioneer infant microbiome. Microbiome 2017;5:1–19.

Primm TP, Lucero CA, Falkinham JO. Health Impacts of Environmental Mycobacteria. Clin Microbiol Rev 2004;17:98–106.

Quast C, Pruesse E, Yilmaz P et al. The SILVA ribosomal RNA gene database project: Improved data processing and web-based tools. Nucleic Acids Res 2013;41:590–6.

Risely A, Waite DW, Ujvari B et al. Active migration is associated with specific and consistent changes to gut microbiota in Calidris shorebirds. J Anim Ecol 2018;87:428–37.

Roto SM, Kwon YM, Ricke SC. Applications of In Ovo technique for the optimal development of the gastrointestinal tract and the potential influence on the establishment of its microbiome in poultry. Front Vet Sci 2016;3:1–13.

Ryan MP, Adley CC. Ralstonia spp.: Emerging global opportunistic pathogens. Eur J Clin Microbiol Infect Dis 2014;33:291–304.

Salter SJ, Cox MJ, Turek EM et al. Reagent and laboratory contamination can critically impact sequence-based microbiome analyses. BMC Biol 2014;12:1–12.

Soler JJ, Martínez-García Á, Rodríguez-Ruano SM et al. Nestedness of hoopoes’ bacterial communities: symbionts from the uropygial gland to the eggshell. Biol J Linn Soc 2016;118:763–73.

Song L, Wu J, Weng K et al. The salmonella effector Hcp modulates infection response, and affects salmonella adhesion and egg contamination incidences in ducks. Front Cell Infect Microbiol 2022;12:1–13.

Stinson LF, Keelan JA, Payne MS. Identification and removal of contaminating microbial DNA from PCR reagents: impact on low-biomass microbiome analyses. Lett Appl Microbiol 2019;68:2–8.

Strandwitz P. Neurotransmitter modulation by the gut microbiota. Brain Res 2018;1693:128–33.

Sun F, Chen J, Liu K et al. The avian gut microbiota: Diversity, influencing factors, and future directions. Front Microbiol 2022;13:1–16.

Sundaram S, Auriemma M, Howard G et al. Application of membrane filtration for removal of diminutive bioburden organisms in pharmaceutical products and processes. PDA J Pharm Sci Technol 1999;53:186–201.

Těšický M, Krajzingrová T, Eliáš J et al. Inter-annual repeatability and age-dependent changes in plasma testosterone levels in a longitudinally monitored free-living passerine bird. Oecologia 2022;198:53–66.

Těšický M, Krajzingrová T, Świderská Z et al. Longitudinal evidence for immunosenescence and inflammaging in free-living great tits. Exp Gerontol 2021;154:111527.

Teyssier A, Lens L, Matthysen E et al. Dynamics of gut microbiota diversity during the early development of an avian host: Evidence from a cross-foster experiment. Front Microbiol 2018;9, DOI: 10.3389/fmicb.2018.01524.

Trevelline BK, MacLeod KJ, Knutie SA et al. In ovo microbial communities: A potential mechanism for the initial acquisition of gut microbiota among oviparous birds and lizards. Biol Lett 2018;14:3–6.

Tsiodras S, Kelesidis T, Kelesidis I et al. Human infections associated with wild birds. J Infect 2008;56:83–98.

Van Veelen HPJ, Salles JF, Tieleman BI. Microbiome assembly of avian eggshells and their potential as transgenerational carriers of maternal microbiota. ISME J 2018;12:1375–88.

Walker RW, Clemente JC, Peter I et al. The prenatal gut microbiome: are we colonized with bacteria in utero? Pediatr Obes 2017;12:3–17.

Wang Q, Garrity GM, Tiedje JM et al. Naïve Bayesian classifier for rapid assignment of rRNA sequences into the new bacterial taxonomy. Appl Environ Microbiol 2007;73:5261–7.

Wickham H. Ggplot2: Elegant Graphics for Data Analysis. Springer-Verlag New York, 2016.

Wright ES. DECIPHER: Harnessing local sequence context to improve protein multiple sequence alignment. BMC Bioinformatics 2015;16:1–14.

