## Supplementary information 1 (SI1) for "Nearly (?) sterile avian egg in a passerine bird"

**List of supplementary material and methods online (SMMO)**

**List of supplementary tables**

**List of supplementary figures**

**Supplementary materials and methods online (SMMO)**

SMMO 1: Selection of bacterial DNA isolation kit

To choose the most suitable DNA isolation kit for bacterial DNA extraction, we initially compared five following kits: High Pure PCR Template Preparation Kit (No.11796828001, Roche Applied Science, Penzberg, Germany), DNeasy PowerSoil Kit (Qiagen, Cat No. 47016, Hilden, Germany), EliGene MTB Isolation (Elisabeth Pharmacon, Cat. No. 90043-50, Brno, Czech Republic), NucleoSpin Tissue kit (Macherey-Nagel Inc., Cat. No. 740952.50, Bethlehem, USA) and QIAamp cador Pathogen Mini Kit (Qiagen, Cat. No. 54104) using trial chicken egg content and gastrointestinal tract (GIT) samples. First, we assessed kit extraction sensitivity. For doing so, we isolated *Enterococcus faecium* (NCIMB 11181, Lactiferm basic 5, Chr. Hansen, Hørsholm, Denmark) diluted both in chicken egg contents and distilled water in the concentration series 10^-8^ – 10^-4^ Colony Forming Unit (CFU). Then we ran qPCR with newly designed universal bacterial qPCR primers (forward: 16Sr_RNA_bact4_F, AAACTCAAAKGAATTGACGGGG, and reverse: 16S_rRNA_bact4_R, TCACRRCACGAGCTGAC, amplification of 172 bp amplicon) and 2x qPCRBIO SyGreen Blue Mix Lo-ROX Kit (PCR Biosystems Cat. No. PB20.15-01, London, England) according to the manual. Based on the highest kit sensitivity (i.e. highest C_p_ value in the lowest CFU concentration; data not shown), we further selected DNeasy PowerSoil Kit and High Pure PCR Template Preparation Kit for further testing. Second, we extracted bacterial DNA from chicken caeca and small intestine using these two kits under different tissue treatments: (1) tissue homogenization in bead tubes in MagNAlyzer (Roche Applied Science, Switzerland), (2) tissue homogenization in tissue lyzer (Retsch MM200, Retsch GmbH, Haan, Germany) steel bead with ca 5 mm in diameter, (3) vortexing with the steel bead (vortex Vortex-Genie2 equipped with MolBio adapter, MolBio Laboratories, Inc.) and (4) tissue vortexing without a bead; in total of 5 samples per kit and treatment). Then using qPCR with the same kit as above, we measured the ratio between 16S rRNA bacterial DNA (the primer as above) and eukaryotic 28S rRNA DNA primers (forward: 28SFwd GACGACCGATTTGCACGTC, reverse: 28SRev GACGACCGATTTGCACGTC, amplifying 62 bp amplicon; Vinkler et al. 2018). During microbial extraction, the isolation of undesirable eukaryotic DNA (assessed e.g. by 28S rRNA DNA qPCR) cannot be avoided but its concentration should be minimised as it could negatively affect PCR amplification of samples with low bacterial biomass. As we expected the higher 16S rRNA/ 28S rRNA DNA concentration ratio, the better, we selected treatment (4) “tissue vortexing without bead“ for subsequent analysis. Finally, to reveal the potential kit isolation bias in taxonomical composition and potential kit-specific contaminations, we sequenced the same chicken egg content (N = 10) and GIT samples (N =10) including isolation negative control and negative control of PCR. We used bacterial universal primers (S-D-Bact-0341-b-S17: CCTACGGGNGGCWGCAG and S-D-Bact-0785-a-A-21: (GACTACHVGGGTATCTAATCC) according to our previously published protocol (Kropáčková et al., 2017). Based on the sequencing results (data are not shown), DNeasy PowerSoil Kit was selected for all subsequent extractions.

SMMO 2: Deviation from the recommended procedure for DNA extraction by DNeasy PowerSoil Kit

Steps according to DNeasy PowerSoil Kit Handbook (Quiagen 2017):

Step 1: For egg content samples (E0-egg), 200 μL homogenized egg content mixed with 200 μL sterile water was added.

Step 2: For E0-egg samples, 200 μl solution C1 instead of 60 μl was added.

Step 17: To concentrate eluted DNA, 55 µl C6 solution instead of 100 μl was added at the center of the white filter membrane.

SMMO 3: In ovo *Enterococcus* and PBS administration procedure

To confirm that bacteria present in the egg content of freshly laid eggs (E0-egg) can be detected with our methods later during the embryonic life, we injected within two days after laying a subset of the eggs (one egg per nest, N = 55) with *Enterococcus faecium* (ref. strain: NCIB 11181 in Lactiferm Basic 5, Chr. Hansen, Hørsholm, Denmark Cat. No.: L-0265). Specifically, we injected the first egg per clutch with 10 µl of *E. faecium* in a concentration of either 10^7^ (high dose; N = 28) or 10^4^ (low dose; N = 27) CFU and the second egg with the 10 µl of PBS (Sigma Aldrich, Cat. No. D5652-50L; N = 53), serving as a control.

In the field, the sterile probiotic powder was always freshly mixed with PCR clean water (Qiagen, Cat. No.: 129114) and kept in cold and dark before application (max 20 min – 1 hour). PBS solution was prepared, aliquoted and kept frozen before application. To prevent contamination, we wore sterile gloves and before the injection, the site of injection was cleaned with 96% EtOH. The injection was done using a sterile insulin syringe Omnican 50 (B Braun, Cat. No. 9151125, Melsungen, Germany) by punctuating the eggshell on a blunt end, which was followed by sealing the aperture with a superglue. After the manipulation, the treated eggs were given back to their nest and together with E13-nat were let incubated until E13 when all eggs were collected. Tissue samples were successfully obtained in a total of 23 E13-Ent and 29 E13-PBS eggs, i.e. 41.1 % and 54.7 % of initial numbers because of embryos´ mortality. Four E13-Ent and two E13-PBS embryos were hatched before the collection (see Figure 1 for a timeline and experimental design scheme).

**Supplementary tables**

Table S1: PCR conditions used for amplification in the first and the second PCR reactions with no-chloroplast amplifying primers (A; protocol 1, P1) and bacterial universal primers (B; protocol 2, P2) and in egg content, embryonic and female faeces sample

PCR reactions were performed in two independent steps. For both protocols, the first PCR was performed in a total volume of 10 µl and the second PCR in a total volume of 20 µl. After the first PCR, the amplified PCR products were diluted 12x in P1 and 24x in P2 before the second PCR was performed. While for P2 identical conditions were applied for both egg content and embryonic samples, for P1 protocols differing in the number of cycles were used. For female feces samples, the only P1 protocol was used. Legend: ^a^ – egg content, ^b^ – embryos, ^c^ – female faeces.

**A)**

| **PCR volumes** |  |  |
| --- | --- | --- |
| **First PCR** | **Volume** |  |
| Primer mix (0.1 µM F + R) | 1.2 µl |  |
| Buffer A | 2 µl |  |
| dNTP | 0.2 µl |  |
| MgCl2 | 0.3 µl |  |
| Kapa Robust polymerase | 0.1 µl |  |
| RNAse free water | 1.2 µl |  |
| DNA | 5 µl |  |
| **Total** | **10 µl** |  |
| **Second PCR** | **Volume** |  |
| barcodes | 4 µl |  |
| Buffer A | 4 µl |  |
| dNTP | 0.4 µl |  |
| MgCl2 | 0.6 µl |  |
| Kapa Robust polymerase | 0.16 µl |  |
| RNAse free water | 8.84 µl |  |
| PCR product (12x diluted) | 2 µl |  |
| **Total** | **20 µl** |  |
| **PCR conditions** |  |  |
|  | **First PCR** | **Second PCR** |
| Initial PCR activation | 95°C/ 3 min | 95°C/ 3 min |
| Denaturation | 95°C/ 30 sec | 95°C/ 15 sec |
| Annealing | 57°C/ 30 sec | 55°C/ 30 sec |
| Extension | 72°C/ 25 sec | 72°C/ 30 sec |
| Number of cycles | 28^a^/ 28^b^ /30^c^ | 24^a^/ 20^b^ /16^c^ |
| Final extension | 72°C/ 1 min | 72°C/ 3 min |

**B)**

| **PCR volumes** |  |  |
| --- | --- | --- |
| **First PCR** | **Volume** | |
| Master mix | 5 µl |  |
| Primer mix (0.1 µM F + R) | 0.4 µl |  |
| RNA free-water | 0 µl |  |
| DNA | 4.6 µl |  |
| **Total** | **10 µl** |  |
| **Second PCR** | **Volume** | |
| Master mix | 10 µl | |
| Adaptors | 4 µl | |
| RNA free-water | 0 µl | |
| PCR product (24x diluted) | 6 µl | |
| **Total** | **20 µl** |  |
| **PCR conditions** |  |  |
|  | **First PCR** | **Second PCR** |
| Initial PCR activation | 95°C/ 3 min | 95°C/ 3 min |
| Denaturation | 95°C/ 30 sec | 95°C/ 30 sec |
| Annealing | 55°C/ 30 sec | 55°C/ 30 sec |
| Extension | 72°C/ 30 sec | 72°C/ 30 sec |
| Number of cycles | 30^a,b^ | 18^a,b^ |
| Final extension | 72°C/ 5 min | 72°C/ 5 min |

Table S2: Number of collected samples and number of samples, sequences and bacterial amplicon sequence variants (ASVs) after applying different bioinformatic filtering steps (Tables A-E)

Filtering step – bioinformatic filtering steps (A-E, see Table S2A for the explanation), No.all.seq – number of all sequences, No.taxa – number of bacterial ASVs, No.all samples – total number of all samples, Numbers of samples with bacterial sequences in given sample type based on different amplification protocols (P1 and P2, see methods for more details) are shown. Samples sequenced only by P1 are labelled by an asterisk. Please note, that number of samples in different sample types for steps A-C is indicated in technical duplicates (i.e. real number of samples * 2). E0-egg – egg content sample at embryonic day 0, E13-nat – E13 intestinal sample from non-manipulated egg, E13-Ent – E13 intestinal sample from *Enterococcus*-treated egg, E13-PBS – E13 intestinal sample from a control PBS-injected egg, F – adult female faecal sample, INC – negative control of isolation (isolation negative control), NTC – negative control of PCR (no template PCR control).

**A) General summary and number of sequences**

| **Filtering step** | **No.all.seq** | **No.taxa** | **No.all samples** | **E0-egg** | **E13-nat** | **E13-prob** | **E13-PBS** | **F*** | **INC** | **NTC** |
| --- | --- | --- | --- | --- | --- | --- | --- | --- | --- | --- |
| Samples collected in the field | – | – | 208 | 52 | 66 | 23 | 29 | 34 | – | – |
| All samples after library preparation before sequencing | – | – | 857 | 104/97 | 132/127 | 46/46 | 58/58 | 63 | 55/44 | 15/12 |
| **(A)** Initial state after sequencing | 1 996 026 | 1382 | 855 | 104/97 | 132/127 | 46/46 | 58/58 | 61 | 55/44 | 15/12 |
| **(B)** Removing samples with low number of sequences (< 50) | 1 994 128 | 1370 | 773 | 90/92 | 128/124 | 43/45 | 52/56 | 54 | 33/42 | 3/11 |
| **(C)** Removing non-consistent ASVs in both duplicates and further samples with (< 50 sequences) | 1 798 405 | 152 | 416 | 47/52 | 66/64 | 23/23 | 29/29 | 30 | 22/23 | 2/6 |
| **(D)** Removing contaminants identified by the statistical method and further samples with (< 50 sequences) | 1 711 376 | 140 | 411 | 47/52 | 66/64 | 23/23 | 29/29 | 30 | 20/23 | 1/4 |
| **(E)** Removing *Ralstonia*/ *Enterococcus* and further samples with (< 50 sequences) | 401 089 | 128 | 191 | 8/37 | 31/44 | 1/2 | 5/7 | 30 | 7/14 | 1/4 |

**B) Total number of sequences**

| Sample type | **A** | **B** | **C** | **D** | **E** |
| --- | --- | --- | --- | --- | --- |
| **E0-egg-P1** | 53700 | 53300 | 45194 | 44850 | 1236 |
| **E0-egg-P2** | 289581 | 289575 | 219254 | 199473 | 25097 |
| **E13-Ent-P1** | 264598 | 264538 | 261495 | 261468 | 82 |
| **E13-Ent-P2** | 325581 | 325572 | 323082 | 322890 | 141 |
| **E13-nat-P1** | 341518 | 341380 | 323342 | 305948 | 45488 |
| **E13-nat-P2** | 215332 | 215293 | 197975 | 184478 | 35414 |
| **E13-PBS-P1** | 40011 | 39809 | 37017 | 36956 | 2787 |
| **E13-PBS-P2** | 24354 | 24316 | 20399 | 19320 | 2841 |
| **F-P1** | 279461 | 279323 | 275280 | 275251 | 272337 |
| **INC-P1** | 13467 | 12814 | 11070 | 10058 | 1123 |
| **INC-P2** | 125299 | 125297 | 75315 | 47724 | 11583 |
| **NTC-P1** | 459 | 270 | 214 | 57 | 57 |
| **NTC-P2** | 22665 | 22641 | 8768 | 2903 | 2903 |

**C) Mean number of sequences per sample**

| Sample type | **A** | **B** | **C** | **D** | **E** |
| --- | --- | --- | --- | --- | --- |
| **E0-egg-P1** | 516.35 | 592.22 | 961.57 | 954.26 | 154.50 |
| **E0-egg-P2** | 2985.37 | 3147.55 | 4216.42 | 3836.02 | 678.30 |
| **E13-Ent-P1** | 5752.13 | 6152.05 | 11369.35 | 11368.17 | 82.00 |
| **E13-Ent-P2** | 7077.85 | 7234.93 | 14047.04 | 14038.70 | 70.50 |
| **E13-nat-P1** | 2587.26 | 2667.03 | 4899.12 | 4635.58 | 1467.35 |
| **E13-nat-P2** | 1695.53 | 1736.23 | 3093.36 | 2882.47 | 804.86 |
| **E13-PBS-P1** | 689.84 | 765.56 | 1276.45 | 1274.34 | 557.40 |
| **E13-PBS-P2** | 419.90 | 434.21 | 703.41 | 666.21 | 405.86 |
| **F-P1** | 4581.33 | 5172.65 | 9176.00 | 9175.03 | 9077.90 |
| **INC-P1** | 244.85 | 388.30 | 503.18 | 502.90 | 160.43 |
| **INC-P2** | 2847.70 | 2983.26 | 3274.57 | 2074.96 | 827.36 |
| **NTC-P1** | 30.60 | 90.00 | 107.00 | 57.00 | 57.00 |
| **NTC-P2** | 1888.75 | 2058.27 | 1461.33 | 725.75 | 725.75 |

**D) Total number of ASV**

| Sample type | **A** | **B** | **C** | **D** | **E** |
| --- | --- | --- | --- | --- | --- |
| **E0.egg.P1** | 190 | 185 | 20 | 16 | 11 |
| **E0.egg.P2** | 294 | 291 | 22 | 17 | 15 |
| **E13.Ent.P1** | 95 | 95 | 16 | 15 | 1 |
| **E13.Ent.P2** | 69 | 68 | 14 | 12 | 3 |
| **E13.nat.P1** | 429 | 429 | 54 | 49 | 36 |
| **E13.nat.P2** | 436 | 435 | 58 | 53 | 50 |
| **E13.PBS.P1** | 117 | 117 | 25 | 23 | 15 |
| **E13.PBS.P2** | 143 | 143 | 18 | 16 | 13 |
| **F.P1** | 80 | 80 | 50 | 49 | 47 |
| **INC.P1** | 87 | 77 | 27 | 19 | 18 |
| **INC.P2** | 152 | 152 | 23 | 14 | 13 |
| **NTC.P1** | 23 | 9 | 6 | 2 | 2 |
| **NTC.P2** | 47 | 47 | 5 | 2 | 2 |

**E) Mean number of ASVs per sample**

| Sample type | **A** | **B** | **C** | **D** | **E** |
| --- | --- | --- | --- | --- | --- |
| **E0-egg-P1** | 4.82 | 5.29 | 1.79 | 1.64 | 2.38 |
| **E0-egg-P2** | 7.71 | 8.08 | 3.12 | 2.29 | 1.65 |
| **E13-Ent-P1** | 6.00 | 6.33 | 3.22 | 3.17 | 1.00 |
| **E13-Ent-P2** | 4.85 | 4.91 | 2.48 | 2.30 | 1.50 |
| **E13-nat-P1** | 10.39 | 10.66 | 3.98 | 3.53 | 4.39 |
| **E13-nat-P2** | 12.23 | 12.49 | 5.53 | 4.34 | 4.36 |
| **E13-PBS-P1** | 5.14 | 5.58 | 2.55 | 2.48 | 3.80 |
| **E13-PBS-P2** | 6.34 | 6.50 | 2.55 | 2.00 | 2.43 |
| **F-P1** | 4.97 | 5.48 | 4.07 | 4.03 | 3.53 |
| **INC-P1** | 3.95 | 5.30 | 3.59 | 2.75 | 3.86 |
| **INC-P2** | 8.89 | 9.26 | 3.57 | 2.00 | 1.71 |
| **NTC-P1** | 2.00 | 3.67 | 3.00 | 2.00 | 2.00 |
| **NTC-P2** | 6.58 | 7.00 | 2.00 | 1.25 | 1.25 |

Table S3: qPCR primer and probe sequences used for the detection of three potentially pathogenic bacteria (*Clostridium*, *Corynebacterium* and *Dietzia*)

F – forward primer, R – reverse primer, P – probe. All probes are in the forward orientation. qPCR efficiency was assessed from serial dilutions of gBlock synthetic standards in the concentration series: 10^9^ – 10^1^ (see Table S4).

| **Gene** | **Target** | **Name** | **Type** | **Sequence (5´- 3´)** | **Length** | **Prod.Length** | **Efficiency** | **Probe orientation** |
| --- | --- | --- | --- | --- | --- | --- | --- | --- |
| *16S rRNA* | *Clostridium* | Clostridium_16S_rRNA-F1 | F | CGGATGATTAAGTGGGAT | 18 | 82 | 1.995 | F |
| *16S rRNA* | *Clostridium* | Clostridium_16S_rRNA-R1 | R | TCTCCTGCACTCTAGATAA | 18 |  |  |  |
| *16S rRNA* | *Clostridium* | Clostridium_16S_rRNA-P1 | P | CAACTTGGGTGCTGCATTCCAA | 22 |  |  |  |
| *16S rRNA* | *Corynebacterium* | Corynebacterium_16SrRNA-F1 | F | GCAGGCGATACGG | 13 | 117 | 1.989 |  |
| *16S rRNA* | *Corynebacterium* | Corynebacterium_16SrRNA-R1 | R | TAACTGCCCAGTAACC | 16 |  |  | F |
| *16S rRNA* | *Corynebacterium* | Corynebacterium_16SrRNA-P1 | P | CATAACTTGAGTACTGTAGGGGTAACT | 27 |  |  |  |
| *16S rRNA* | *Dietzia* | Dietzia_16SrRNA-F1 | F | CCTGTAGTACTCAAGTCTG | 19 | 67 | 1.835 | F |
| *16S rRNA* | *Dietzia* | Dietzia_16SrRNA-R1 | R | TCGTCCGTGAAAACTC | 16 |  |  |  |
| *16S rRNA* | *Dietzia* | Dietzia_16SrRNA-P1 | P | CGTATCGCCCGCAAGCTC | 18 |  |  |  |

Table S4: Synthetic gBlock DNA standards for qPCR TaqMan assays used for detection of three potentially pathogenic bacteria (*Clostridium*, *Corynebacterium* and *Dietzia*)

| **Gene** |  | **Target** | **Standard name** | **Length [bp]** | **Standard sequence (5'- 3')** |
| --- | --- | --- | --- | --- | --- |
| *16S rRNA* |  | *Clostridium* | 16SrRNAClostridium-S1 | 150 | TTATCCGGATTTACTGGGCGTAAAGGGAGCGTAGGCGGATGATTAAGTGGGATGTGAAATACCCGGGCTCAACTTGGGTGCTGCATTCCAAACTGGTTATCTAGAGTGCAGGAGAGGAGAGTGGAATTCCTAGTGTAGCGGTGAAATGCG |
| *16S rRNA* |  | *Corynebacterium* | 16SrRNACorynebacterium-S1 | 150 | GCTTAACTCCGGGCGTGCAGGCGATACGGGCATAACTTGAGTACTGTAGGGGTAACTGGAATTCCTGGTGTAGCGGTGAAATGCGCAGATATCAGGAGGAACACCGATGGCGAAGGCAGGTTACTGGGCAGTTACTGACGCTGAGGAGCG |
| *16S rRNA* |  | *Dietzia* | 16SrRNADietzia-S1 | 150 | TATCTGCGCATTTCACCGCTACACCAGGAATTCCAGTCTCCCCTGTAGTACTCAAGTCTGCCCGTATCGCCCGCAAGCTCGGAGTTAAGCTCCGAGTTTTCACGGACGACGCGACAAACCGCCTACGAGCTCTTTACGCCCAGTAATTCC |

Table S5: Pre-amplification PCR and qPCR conditions used for the detection of three potentially pathogenic bacteria (*Clostridium*, *Corynebacterium,* and *Dietzia*)

Pre-amplification was performed with universal bacterial primers (S-D-Bact-0341-b-S17: CCTACGGGNGGCWGCAG and S-D-Bact-0785-a-A-21: GACTACHVGGGTATCTAATCC) using chemistry Platinum SuperFi PCR I Master Mix (ThermoFisher Scientific, Cat. No. 12351, Waltham, Massachusetts, USA) in a total volume of 12 µl. After pre-amplification, PCR products were diluted 3x in spike water. Bacteria-specific qPCRs were then performed in a total volume of 6 µl with specific newly designed primers and probes (see Table S3 for more details) using Luna Universal Probe qPCR Master Mix (New England Biolabs, Inc., No. E3006, Ipswich, Massachusetts, USA) with identical qPCR conditions for all three assays.

| **PCR volumes** |  |
| --- | --- |
| **Pre-amplification PCR** | **Volume** |
| Master mix | 6 µl |
| Primer mix (5 µM F + R) | 1.2 µl |
| Spike water | 1.2 µl |
| DNA | 3.6 µl |
| **Total** | **12 µl** |
| **DNA Probe-based qPCR** | **Volume** |
| Master mix | 3 µl |
| OTU-specific primer mix (5 µM F + R) | 0.72 µl |
| OTU-specific probe (10 µM) | 0.12 µl |
| Spike water | 1.16 µl |
| Pre-amplified PCR product (3x diluted) | 1 µl |
| **Total** | **6 µl** |

| **PCR conditions** |  |
| --- | --- |
| **Pre-amplification PCR** | **Conditions** |
| Initial PCR activation | 95°C/ 3 min |
| Denaturation | 95°C/ 30 sec |
| Annealing | 55°C/ 30 sec |
| Extension | 72°C/ 30 sec |
| Number of cycles | 30 |
| Final extension | 72°C/ 5 min |
| **TaqMan qPCR** | **Conditions** |
| Initial denaturation | 95°C/ 1 min |
| Extension | 95°C/ 20 sec |
|  | 60°C/ 1 min |
| Number of cycles | 45 |
| Cooling | 40°C/ 10 sec |

Table S6: Protest analysis of consistency between technical PCR duplicates (i.e. procrustean analysis running on Bray-Curtis dissimilarities)

Results for different amplification protocols (P1 and P2, see methods for more details) are shown. E0-egg – egg content sample at embryonic day 0, E13-nat – E13 intestinal sample from non-manipulated egg, E13-Ent – E13 intestinal sample from *Enterococcus*-treated egg, E13-PBS – E13 intestinal sample from a control PBS-injected egg, INC – negative control of isolation (i.e. isolation negative control).

| **Sample.type.protocol** | **Significance** | **Sum.of.Squares** | **Correlation** |
| --- | --- | --- | --- |
| **F_P1** | 0.001 | 0.030 | 0.985 |
| **E0-egg_P1** | 0.067 | 0.699 | 0.549 |
| **E0-egg_P2** | 0.001 | 0.466 | 0.730 |
| **E13-Ent_P1** | 0.001 | 0.027 | 0.987 |
| **E13-Ent_P2** | 0.001 | 0.108 | 0.944 |
| **E13_nat_P1** | 0.001 | 0.136 | 0.930 |
| **E13_nat_P2** | 0.001 | 0.178 | 0.906 |
| **E13_PBS_P1** | 0.001 | 0.205 | 0.891 |
| **E13_PBS_P2** | 0.001 | 0.362 | 0.799 |
| **INC_P1** | 0.001 | 0.084 | 0.957 |
| **INC_P2** | 0.595 | 0.452 | 0.740 |

Table S7: Full taxonomy assignment of identified putatively contaminating amplicon sequence variant (ASV) by Decontam analysis (Davis et al. 2018) in (A) egg content and embryonic samples and (B) female faecal samples of the great tit.

| **(A)** |  |  |  |  |  |  |
| --- | --- | --- | --- | --- | --- | --- |
| **ASV_number** | **Phylum** | **Class** | **Order** | **Family** | **Genus** | **Species** |
| OTU_9417 | Proteobacteria | Gammaproteobacteria | Enterobacterales | Yersiniaceae | <NA> | <NA> |
| OTU_10099 | Proteobacteria | Gammaproteobacteria | Burkholderiales | Methylophilaceae | *Methylotenera* | *versatilis* |
| OTU_10212 | Proteobacteria | Gammaproteobacteria | Burkholderiales | Comamonadaceae | *Curvibacter* | *gracilis* |
| OTU_10259 | Proteobacteria | Gammaproteobacteria | Burkholderiales | Comamonadaceae | *Curvibacter* | <NA> |
| OTU_10512 | Proteobacteria | Alphaproteobacteria | Sphingomonadales | Sphingomonadaceae | *Sphingomonas* | <NA> |
| OTU_10641 | Proteobacteria | Alphaproteobacteria | Rhizobiales | Beijerinckiaceae | *Methylobacterium-Methylorubrum* | <NA> |
| OTU_10819 | Proteobacteria | Alphaproteobacteria | Rhodobacterales | Rhodobacteraceae | *Paracoccus* | <NA> |
| OTU_10987 | Actinobacteriota | Actinobacteria | Corynebacteriales | Nocardiaceae | *Rhodococcus* | <NA> |
| OTU_11163 | Actinobacteriota | Actinobacteria | Micrococcales | Micrococcaceae | *Kocuria* | <NA> |
| OTU_12143 | Firmicutes | Bacilli | Staphylococcales | Staphylococcaceae | *Staphylococcus* | <NA> |
| OTU_12192 | Firmicutes | Bacilli | Staphylococcales | Staphylococcaceae | *Staphylococcus* | <NA> |
| OTU_12277 | Firmicutes | Bacilli | Lactobacillales | Streptococcaceae | *Streptococcus* | <NA> |
| **(B)** |  |  |  |  |  |  |
| **ASV_number** | **Phylum** | **Class** | **Order** | **Family** | **Genus** | **Species** |
| OTU_11070 | Actinobacteriota | Actinobacteria | Corynebacteriales | Mycobacteriaceae | *Mycobacterium* | <NA> |
| OTU_11235 | Actinobacteriota | Actinobacteria | Propionibacteriales | Propionibacteriaceae | *Cutibacterium* | <NA> |

Table S8: Comparison of positivity for *Corynebacterium, Clostridium* and *Dietzia* between specific 16S rRNA qPCR assays and 16S rRNA Next-Generation-Sequencing (NGS) metabarcoding.

E0-egg – egg content sample at embryonic day 0, E13-nat – E13 intestinal sample from non-manipulated egg, E13-Ent – E13 intestinal sample from *Enterococcus*-treated egg, E13-PBS – E13 intestinal sample from a control PBS-injected egg (no *Corynebacterium* and *Clostridium* qPCR assays were run for E13-Ent and E13-PBS), F – adult female faeces sample (number of samples is qPCR assays is lower because of lack of DNA in two samples after the metabarcoding), INC – negative control of isolation (isolation negative control), NTC_qPCR – negative control of PCR (i.e no template qPCR control), NTC_pre-amp – no template PCR control of preamplification step, NTC – negative control of PCR (no template PCR control). Please, note that different biological sample types were analysed in three following batches: (i) E0-egg, (ii) E13-nat, E13-Ent, E13-PBS and (iii) F to prevent cross-contamination between different sample types and, therefore, they have their negative controls. For qPCR results, only samples verified by independent qPCR with pre-amplification step are considered positive (see methods). For NGS data, positivity is assessed only for identical amplicon sequence variants (ASV) for which we designed specific qPCR assays. Samples are considered positive when given ASV was found at least by one amplification protocol (P1 or P2; please see methods) after merging results from independent technical duplicates. Overlap indicates the number of positive samples and their proportion for given ASV both from NGS metabarcoding and specific TaqMan PCR assays.

|  |  | **qPCR** |  |  | **NGS** |  |  |  |
| --- | --- | --- | --- | --- | --- | --- | --- | --- |
| **Biological sample type** | | **INC** | **NTC_qPCR** | **NTC_pre_amplif** | **Biological sample type** | **INC** | **NTC** | **Overlap %** |
| ***Corynebacterium*** |  |  |  |  |  |  |  |  |
| **E0-egg** | 0/52 (0 %) | 0/7 (0 %) | 0/3 (0 %) | 0/9 (0 %) | 2/52 (3.85 %) | 0/7 (0 %) | 0/3 (0 %) | 0/2 (0 %) |
| **E13-nat** | 0/66 (0 %) | 0/16 (0 %) | 0/3 (0 %) | 0/11 (0 %) | 0/66 (0 %) | 0/16 (0 %) | 0/3 (0 %) | 0 % |
| **E13-Ent** | – | – | – | – | 0/23 (0 %) | 0/1 (0 %) | 0/1 (0 %) | 0 % |
| **E13-PBS** | – | – | – | – | 0/29 (0 %) | 0/2 (0 %) | 0/2 (0 %) | 0 % |
| **F** | 8/32 (25.00 %) | 0/3 (0 %) | 0/3 (0 %) | 0/3 (0 %) | 0/34 (0 %) | 0/3 (0 %) | 0/4 (0%) | 0% |
| ***Clostridium*** |  |  |  |  |  |  |  |  |
| **E0-egg** | 0/52 (0 %) | 0/7 (0 %) | 0/3 (0 %) | 0/9 (0 %) | 1/52 (1.92 %) | 0/7 (0 %) | 0/3 (0 %) | 0 % |
| **E13-nat** | 0/66 (0 %) | 0/16 (0 %) | 0/3 (0 %) | 0/9 (0 %) | 0/66 (0 %) | 0/16 (0 %) | 0/3 (0 %) | 0/2 (0 %) |
| **E13-Ent** | – | – | – | – | 0/23 (0 %) | 0/1 (0 %) | 0/1 (0 %) | 0 % |
| **E13-PBS** | – | – | – | – | 0/29 (0 %) | 0/2 (0 %) | 0/2 (0 %) | 0 % |
| **F** | 29/32 (90.63% ) | 1/3 (33.3%) | 0/4 (0 %) | 1/3 (33.3%) | 12/34 (35.29 %) | 0 (0 %) | 0 (0%) | 11/12 (91.66) % |
| ***Dietzia*** |  |  |  |  |  |  |  |  |
| **E0-egg** | 4/52 (7.69 %) | 1/7 (14.29 %) | 0/3 (0 %) | 0/9 (0 %) | 1/52 (1.92 %) | 0/7 (0 %) | 0/3 (0 %) | 1/1 (100 %) |
| **E13-nat** | 2/66 (3.03 %) | 3/16 (18.75 %) | 0/6 (0 %) | 0/18 (0 %) | 2/66 (3.03 %) | 0/16 (0 %) | 0/3 (0 %) | 0/2 (0 %) |
| **E13-Ent** | 0/23 (0 %) | 0/1 (0 %) | 0/1 (0 %) | 0/2 (0 %) | 0/23 (0 %) | 0/1 (0 %) | 0/1 (0 %) | 0 % |
| **E13-PBS** | 1/29 (3.45 %) | 0/2 (0 %) | 0/2 (0 %) | 0/3 (0 %) | 0/29 (0 %) | 0/2 (0 %) | 0/2 (0 %) | 0 % |
| **F** | 29/32 (90.63 %) | 1/3 (33 %) | 0/4 (0 %) | 0/3 (0 %) | 0/34 (0 %) | 0/3 (0 %) | 0/4 (0%) | 0% |

**Supplementary figures**

Figure S1: Proportion of contaminating amplicon sequence variants (ASVs) in different sample types assed by Decontam analysis (Davis et al. 2018) separately in (a) egg content and embryonic samples and (b) in adult females.

Results for different amplification protocols (i.e. P1 and P2; see methods for more details) are shown. Putative contaminating ASVs are mostly present in NTC and INC and are more frequent in samples prepared by P2 protocol (use of more universal primers with DNA polymerase with proofreading activity) than by P1 protocol. E0-egg – egg content sample at embryonic day 0, E13-nat – E13 intestinal sample from non-manipulated egg, E13-Ent– E13 intestinal sample from *Enterococcus*-treated egg, E13-PBS – E13 intestinal sample from a control PBS-injected egg. Negative control of isolation (INC) and PCR (NTC) are also shown.

.

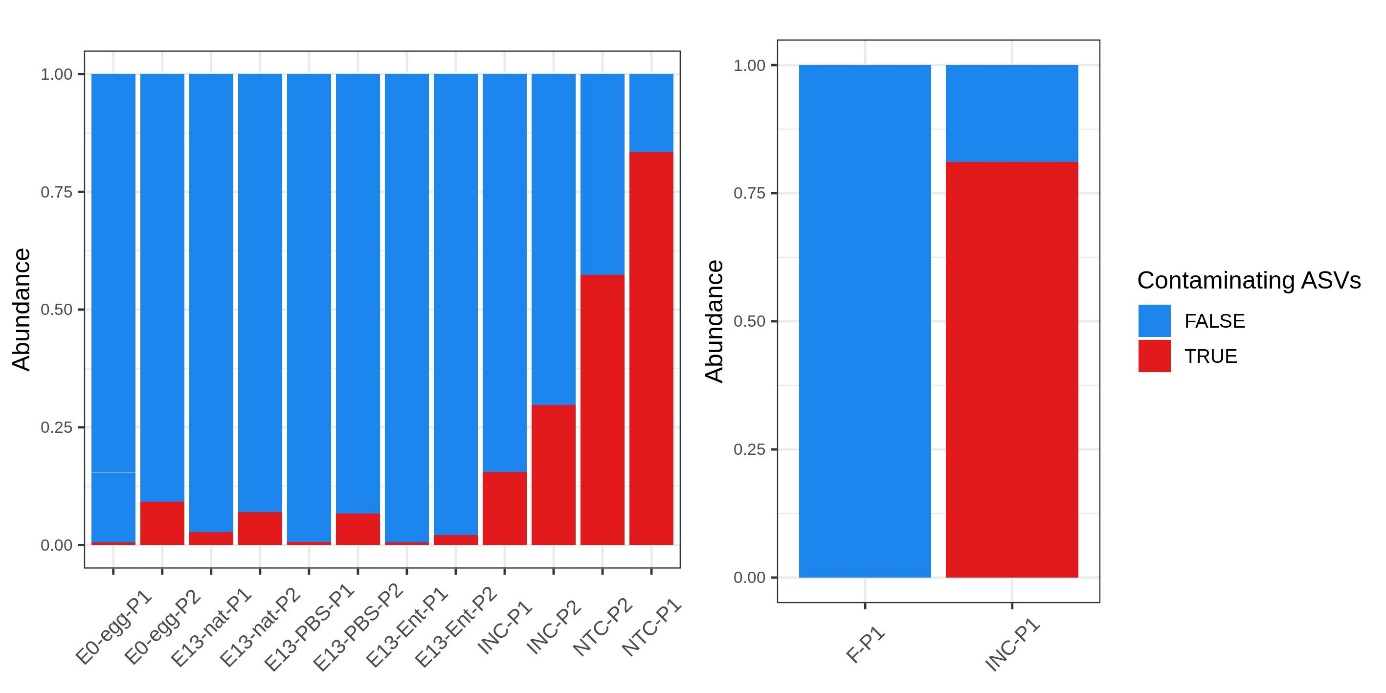

Figure S2: Number of paired read sequences in each sample for different bacterial genera in great tit samples: (A) including potentially contaminating *Ralstonia* and *Enterococcus* and (B) after excluding *Ralstonia* and *Enterococcus*

Only ten most abundant genera are shown. Less abundant genera are included in the category ‘Others’. Results are shown for different amplification protocols (i.e. P1 and P2; see methods for more details). E0-egg – egg content sample at embryonic day 0, E13-nat – E13 intestinal sample from non-manipulated egg, E13-Ent – E13 intestinal sample from *Enterococcus*-treated egg, E13-PBS – E13 intestinal sample from a control PBS-injected egg. Negative control of isolation (INC) and PCR (NTC) are also shown. F – adult female faecal sample.

**(A)**

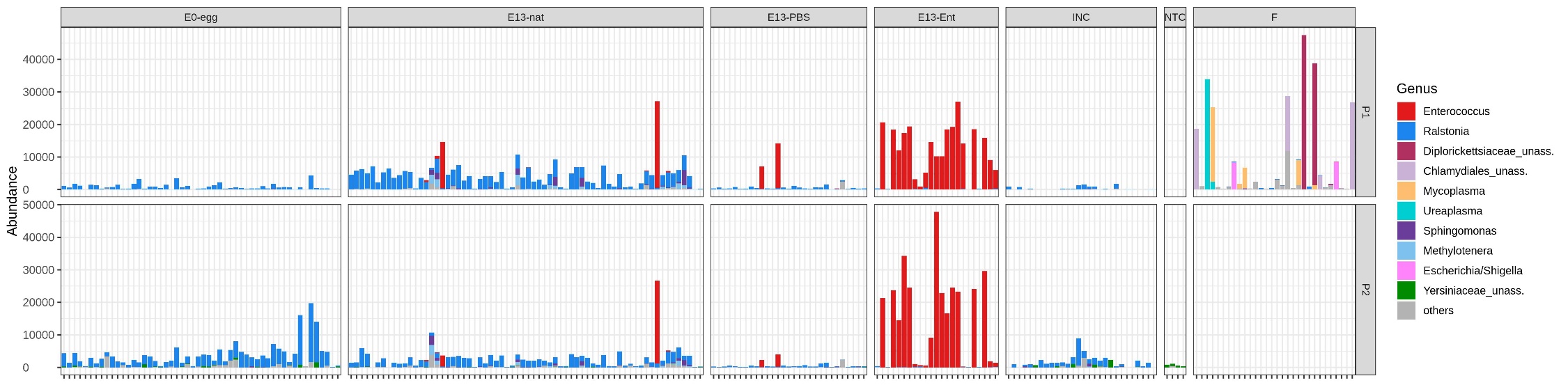

**(B)**

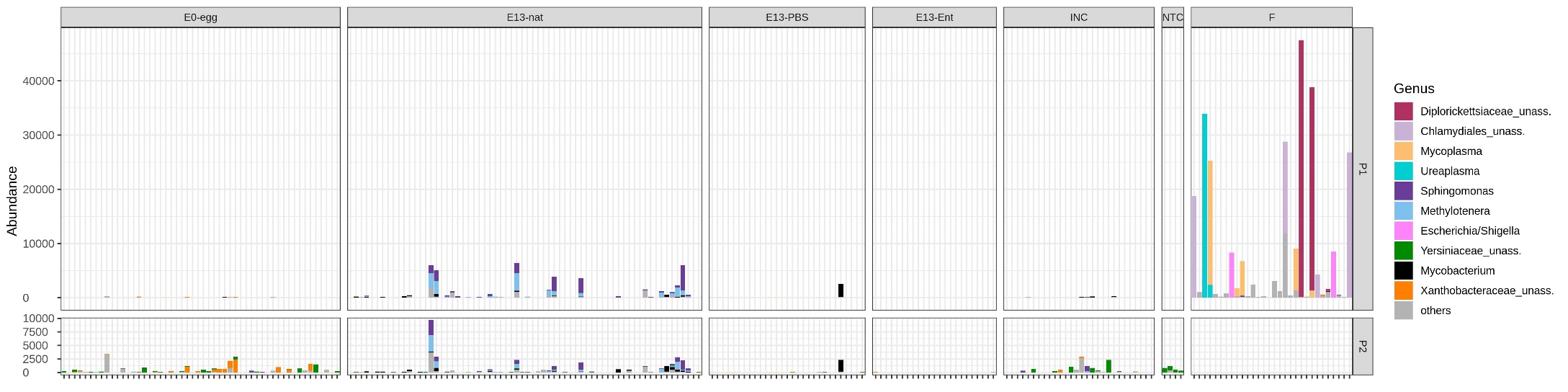

Figure S3: Relative abundance of bacterial phyla (A) and (B) genera in great tit samples.

Only the most abundant phyla (relative abundance > 0.5%) and genera (relative abundance > 1%) are shown. Less abundant phyla and genera are included in the category ‘Others’. Results are shown for different amplification protocols (i.e. P1 and P2; see methods for more details) and including potentially contaminating *Ralstonia* and *Enterococcus*. E0-egg – egg content sample at embryonic day 0, E13-nat – E13 intestinal sample from non-manipulated egg, E13-Ent – E13 intestinal sample from *Enterococcus*-treated egg, E13-PBS – E13 intestinal sample from a control PBS-injected egg. Negative control of isolation (INC) and PCR (NTC) are also shown. F – adult female faecal sample (*amplified only using P1 protocol). For a more detailed taxaplot of female faecal samples, please see Figure S6, SI1.

**(A)**

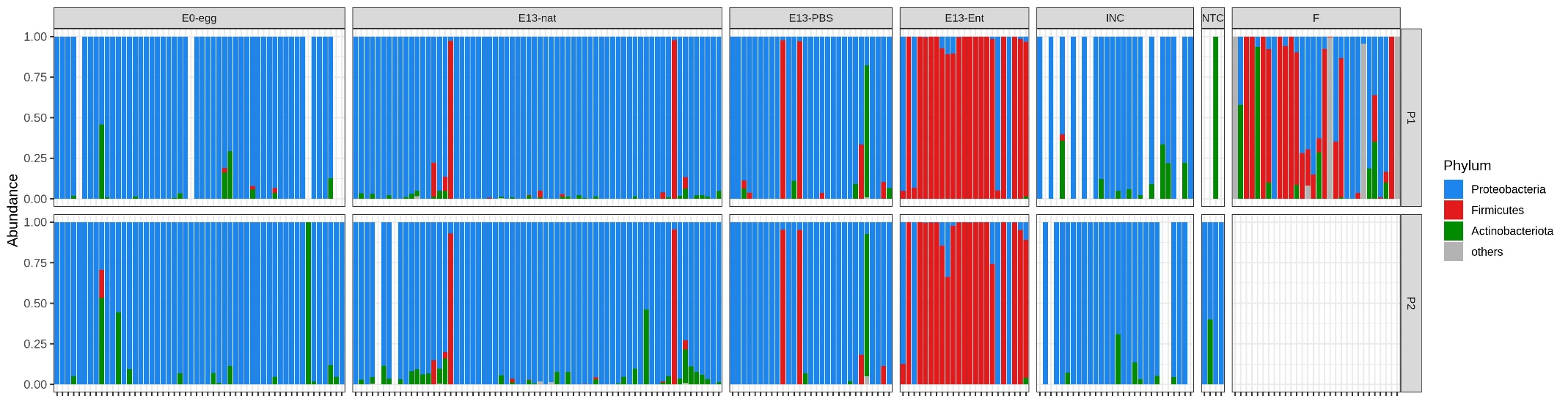

**(B)**

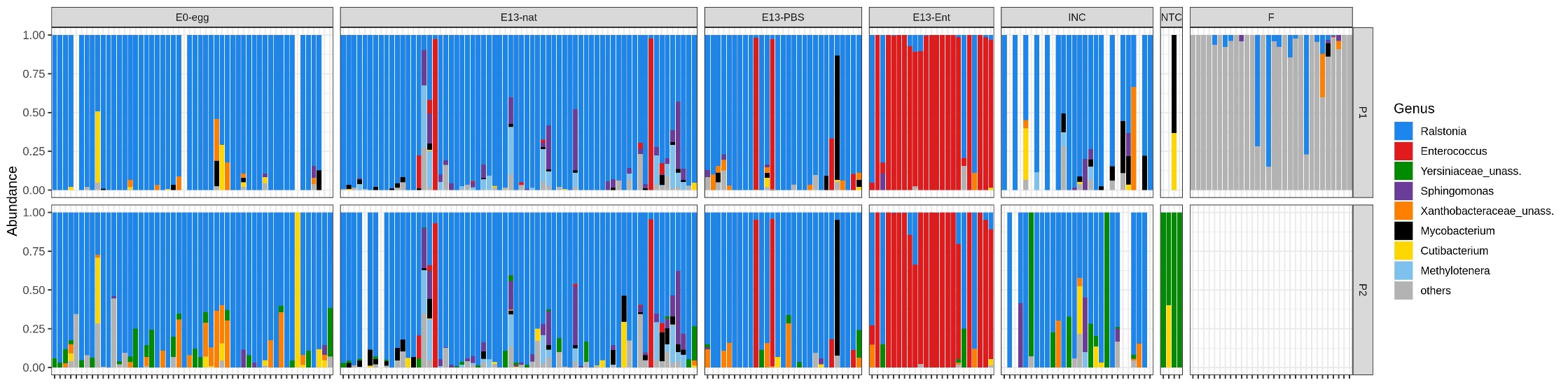

Figure S4: Relative abundances of bacterial taxa after exclusion potentially contaminating *Ralstonia* and cross-contaminating *Enterococcus* sequences for protocol 2 (P2) in great tit egg and embryonic samples

Only the most abundant genera (relative abundance > 1%) are shown. Less abundant genera are included in the category ‘Others’. Only results for P2 are shown (see methods for more details and Figure 4B for comparison with P1). E0-egg – egg content sample at embryonic day 0, E13-nat – E13 intestinal sample from non-manipulated egg, E13-Ent – E13 intestinal sample from a *Enterococcus*-treated egg, E13-PBS – E13 intestinal sample from a control PBS-injected egg.

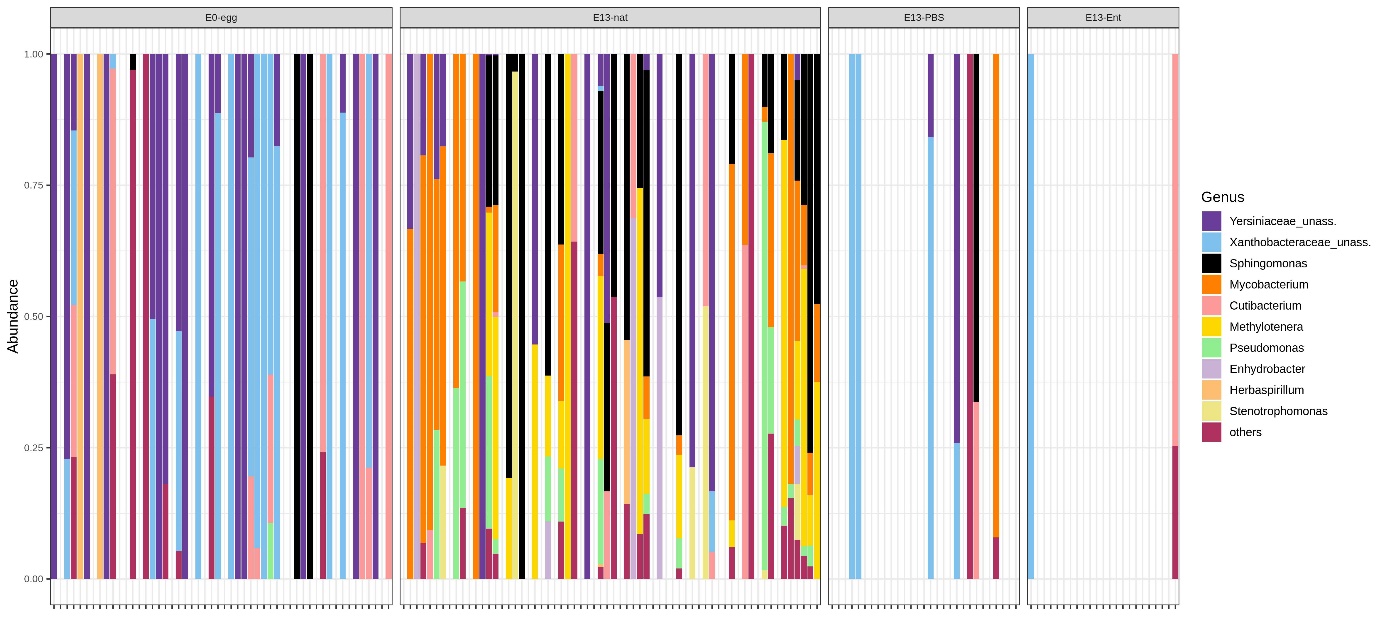

Figure S5: Overall relative abundance of bacterial phyla (a) and (b) genera in the great tit with the summed proportions of all samples per category

Only the most abundant phyla (relative abundance > 0.5%) and genera (relative abundance > 1%) are shown. Less abundant phyla and genera are included in the category ‘Others’. Results are shown for different amplification protocols (i.e. P1 and P2; see methods for more details) and including potentially contaminating *Ralstonia* and *Enterococcus*. E0-egg – egg content sample at embryonic day 0, E13-nat – E13 intestinal sample from non-manipulated egg, E13-Ent – E13 intestinal sample from *Enterococcus*-treated egg, E13-PBS – E13 intestinal sample from a control PBS-injected egg. Negative control of isolation (INC) and PCR (NTC) are also shown.

**(A)**

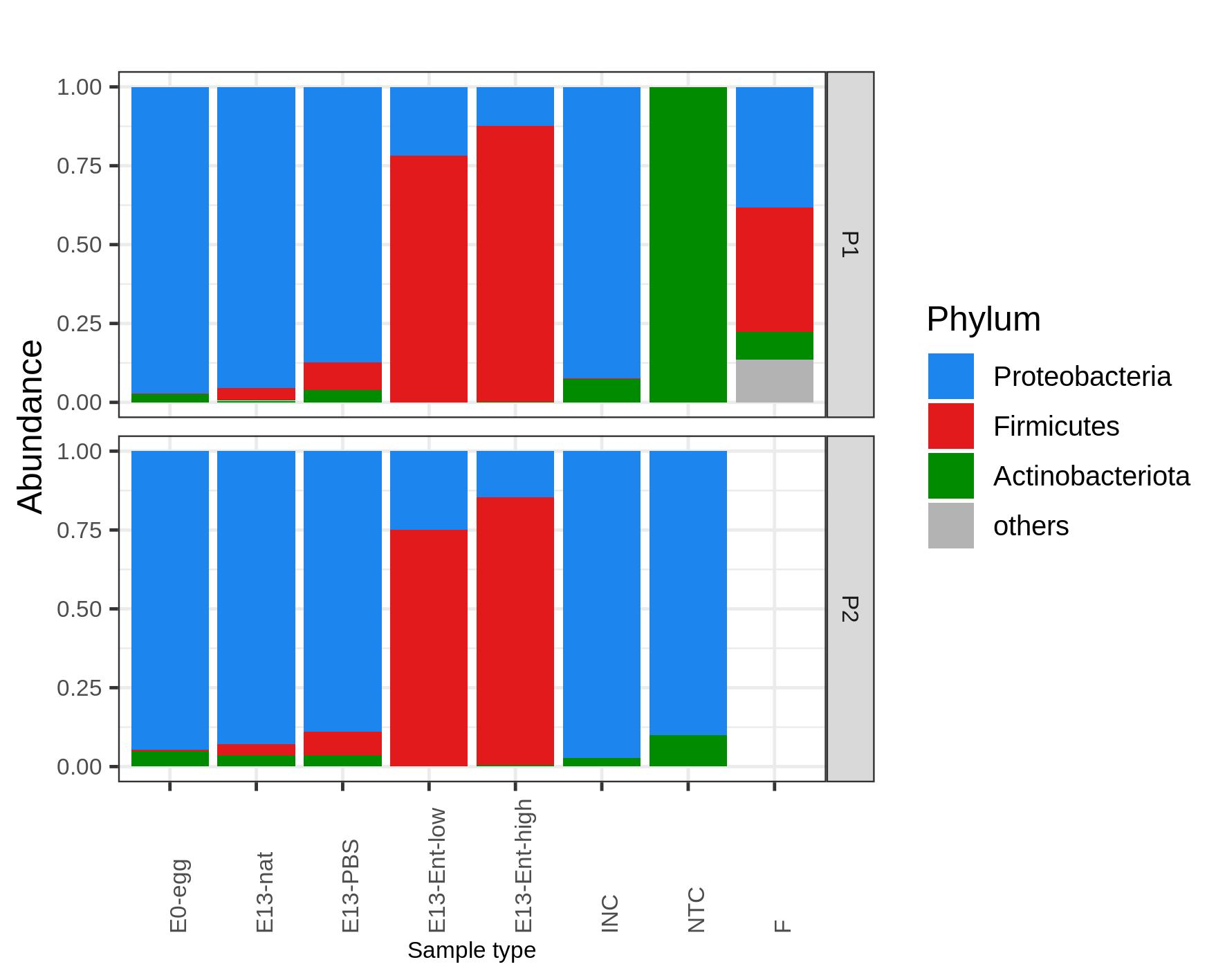

**(B)**

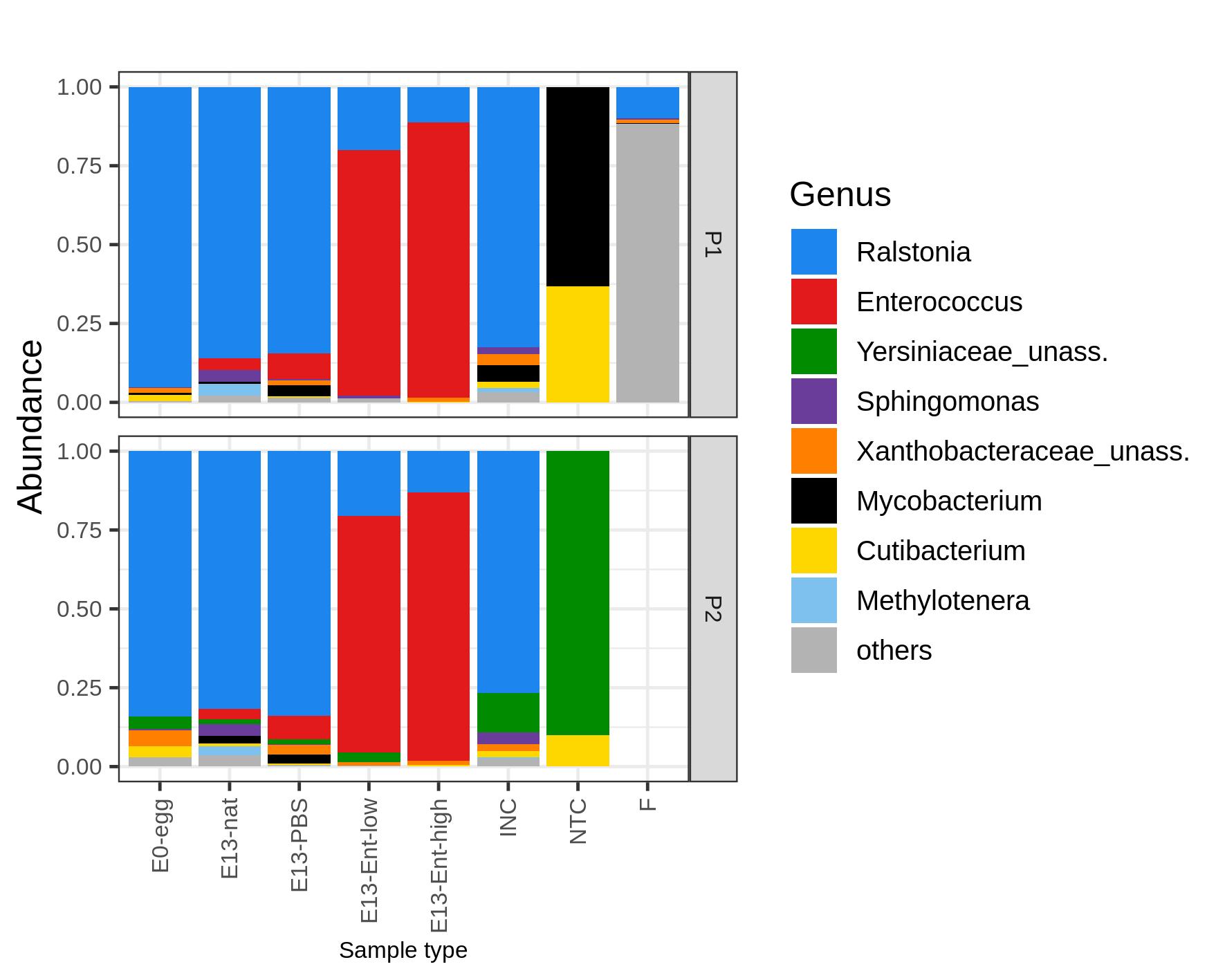

Figure S6: Relative abundance of bacterial phyla (A) and (B) genera in great tit female faecal samples.

Only the most abundant phyla (relative abundance > 0.5%) and genera (relative abundance > 1%) are shown. Results are shown only for amplification protocol 1 (P1; please, see methods for more details).

**(A)**

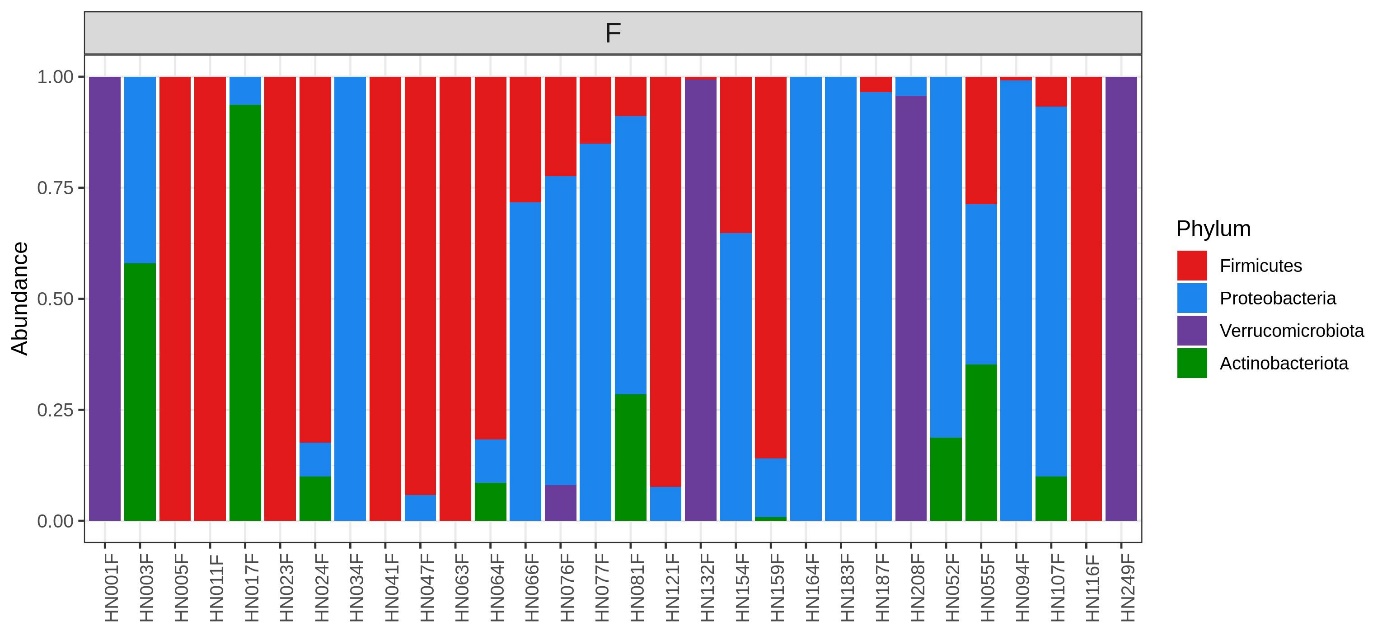

**(B)**

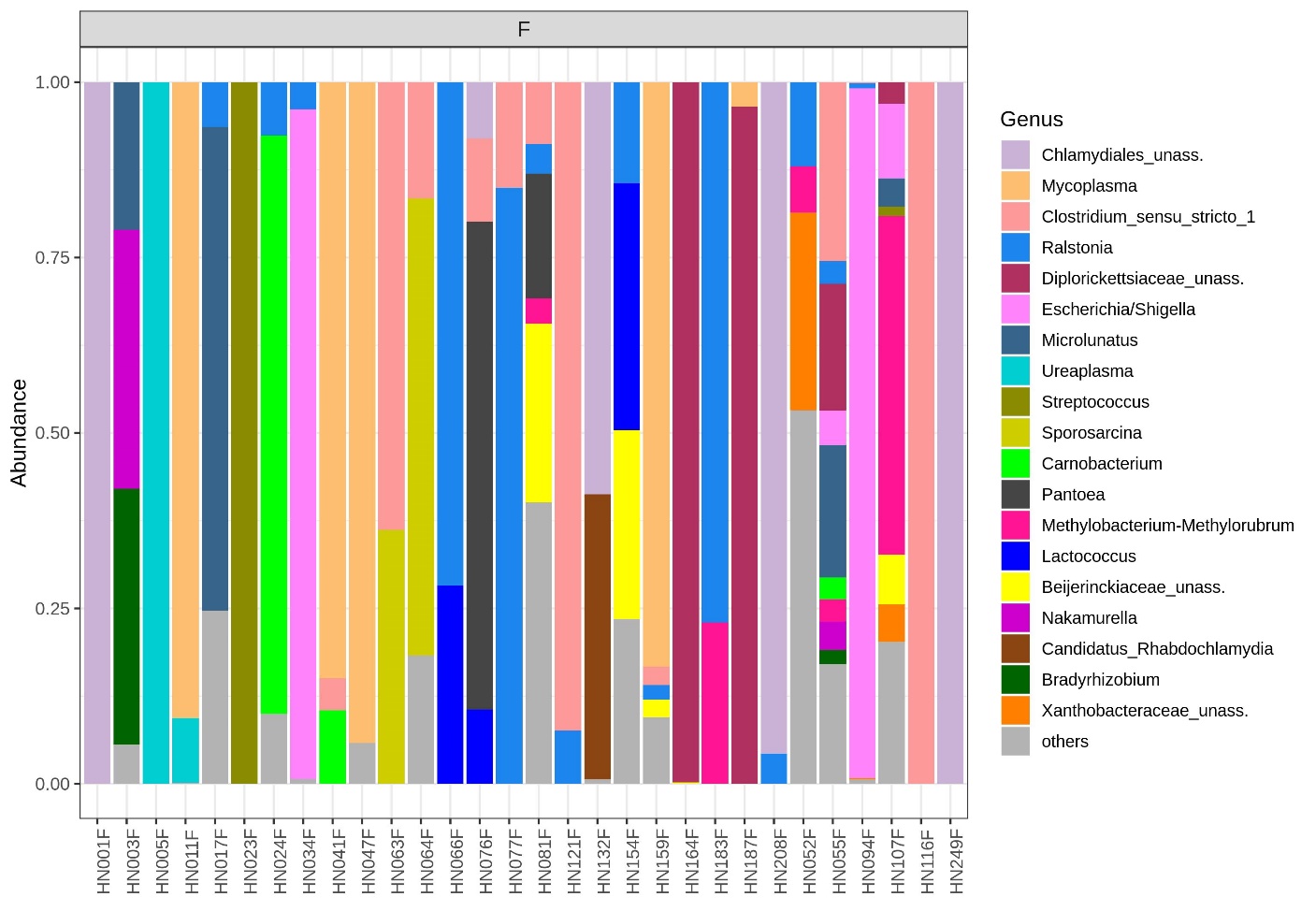

Figure S7: Venn diagrams showing numbers of amplicon sequence variants (ASVs) and genera shared between different biological sample types.

The percentage of sequences belonging to these ASVs/ genera out of all sequences before exclusion of *Ralstonia* and *Enterococcus* is also given. Venn diagrams are presented for protocol 1 (P1) at ASV (A) or genus level (B), for protocol 2 (P2) at ASV (C) or genus level (D) and at ASV (E) or genera level (F) for the combined results from both protocols. E0-egg – egg content sample at embryonic day 0, E13-nat – E13 intestinal sample from non-manipulated egg, E13-Ent – E13 intestinal sample from *Enterococcus*-treated egg, E13-PBS – E13 intestinal sample from a control PBS-injected egg, F – female faecal samples (* female samples were amplified only by P1).

**(A)**

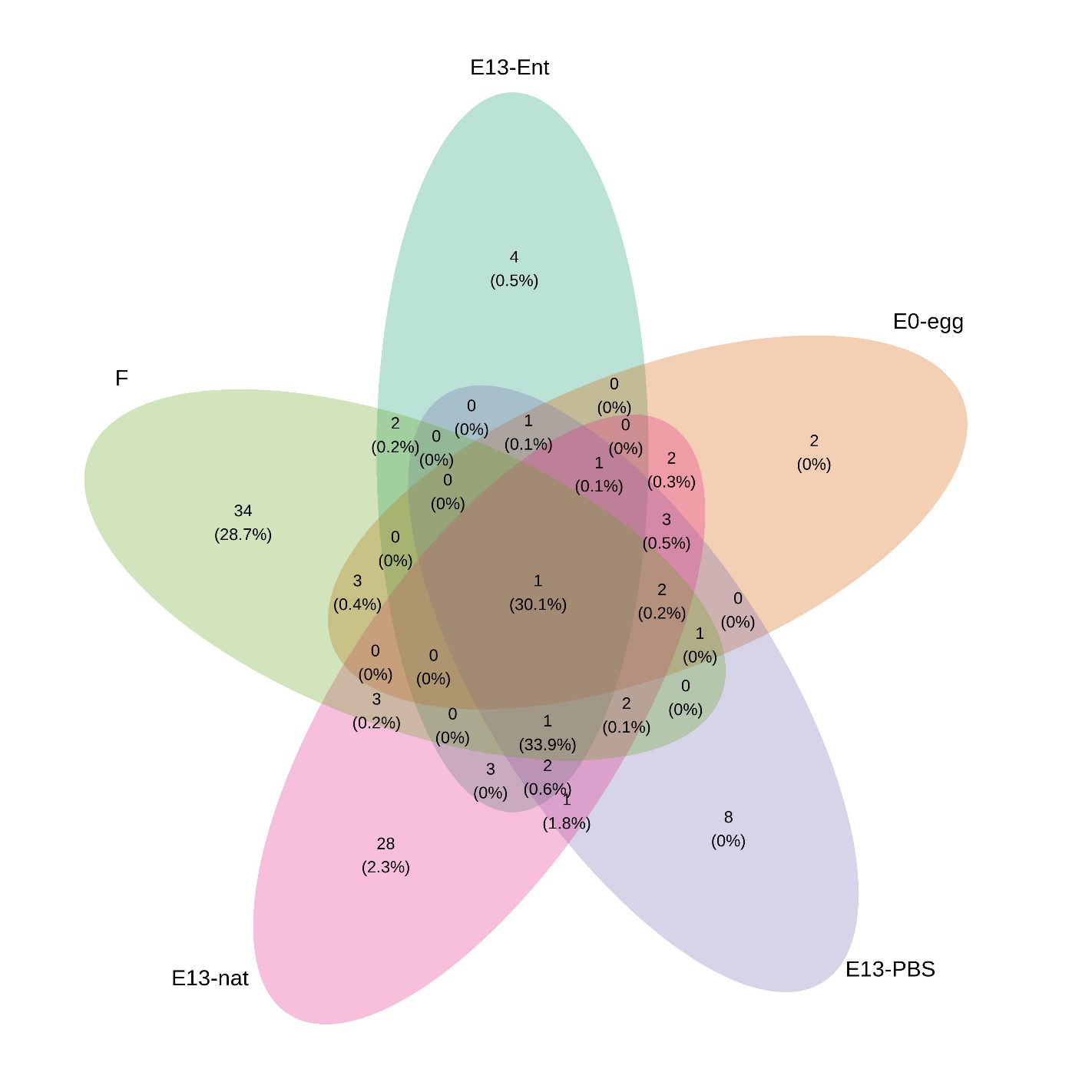

**(B)**

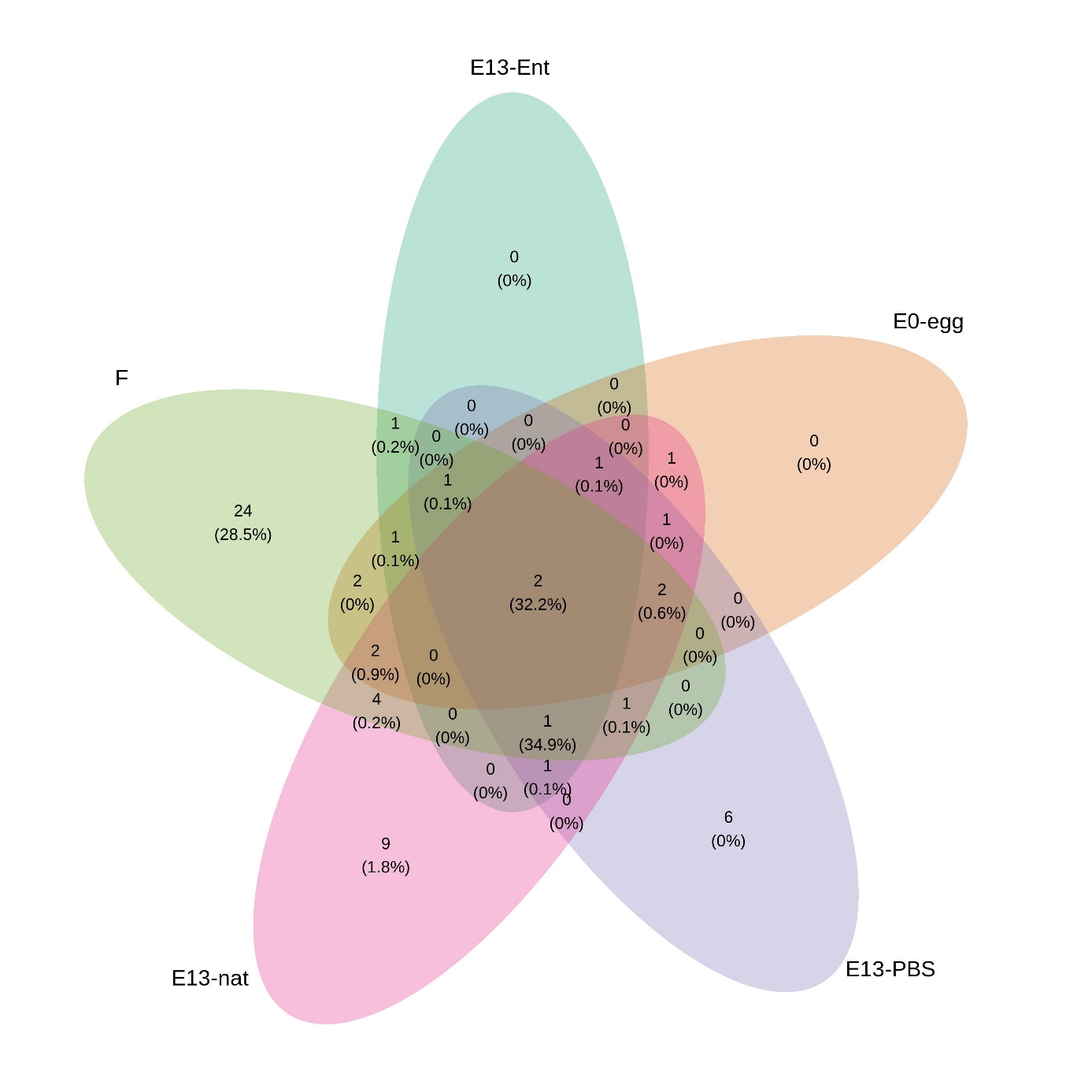

**(C)**

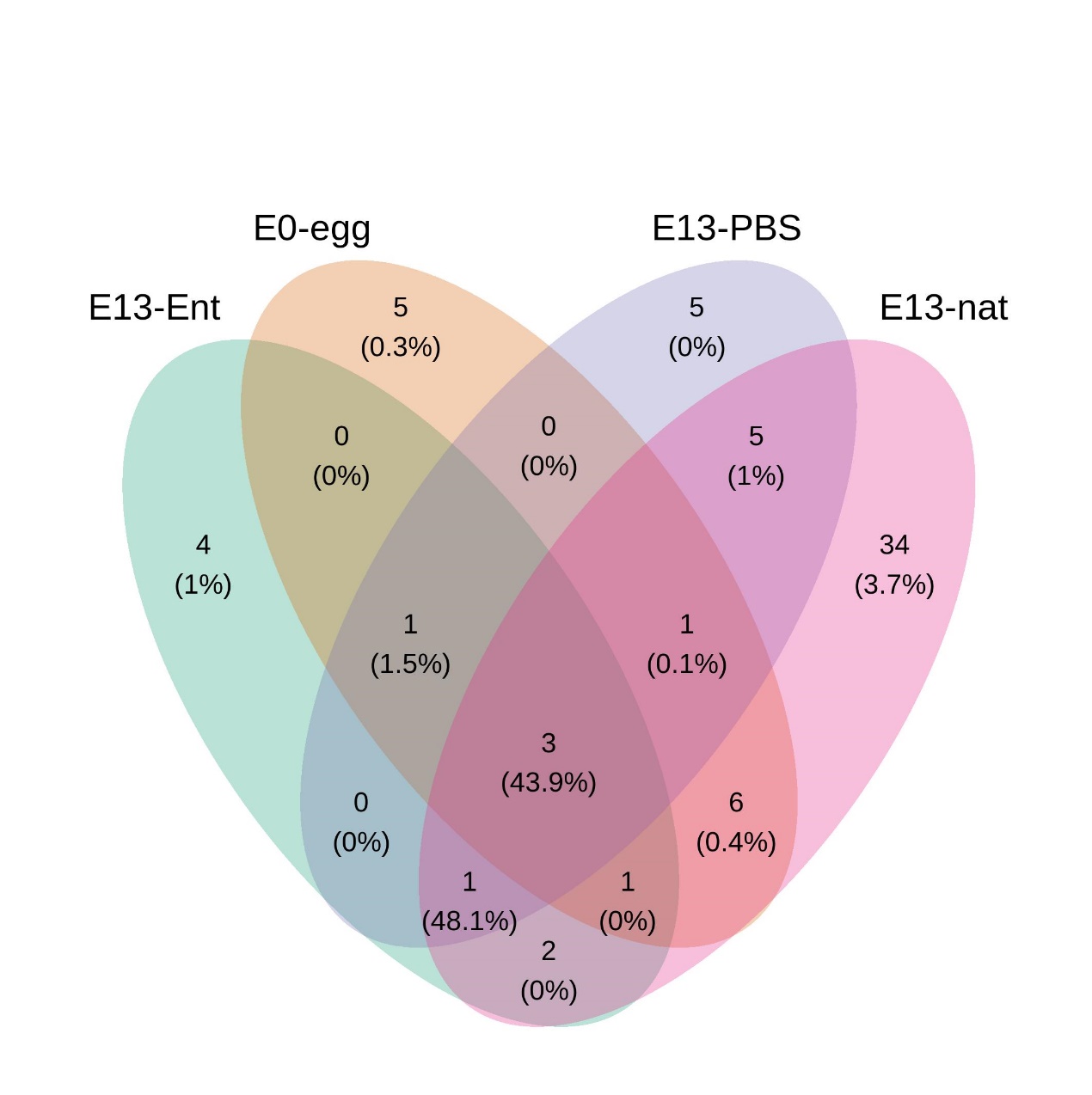

**(D)**

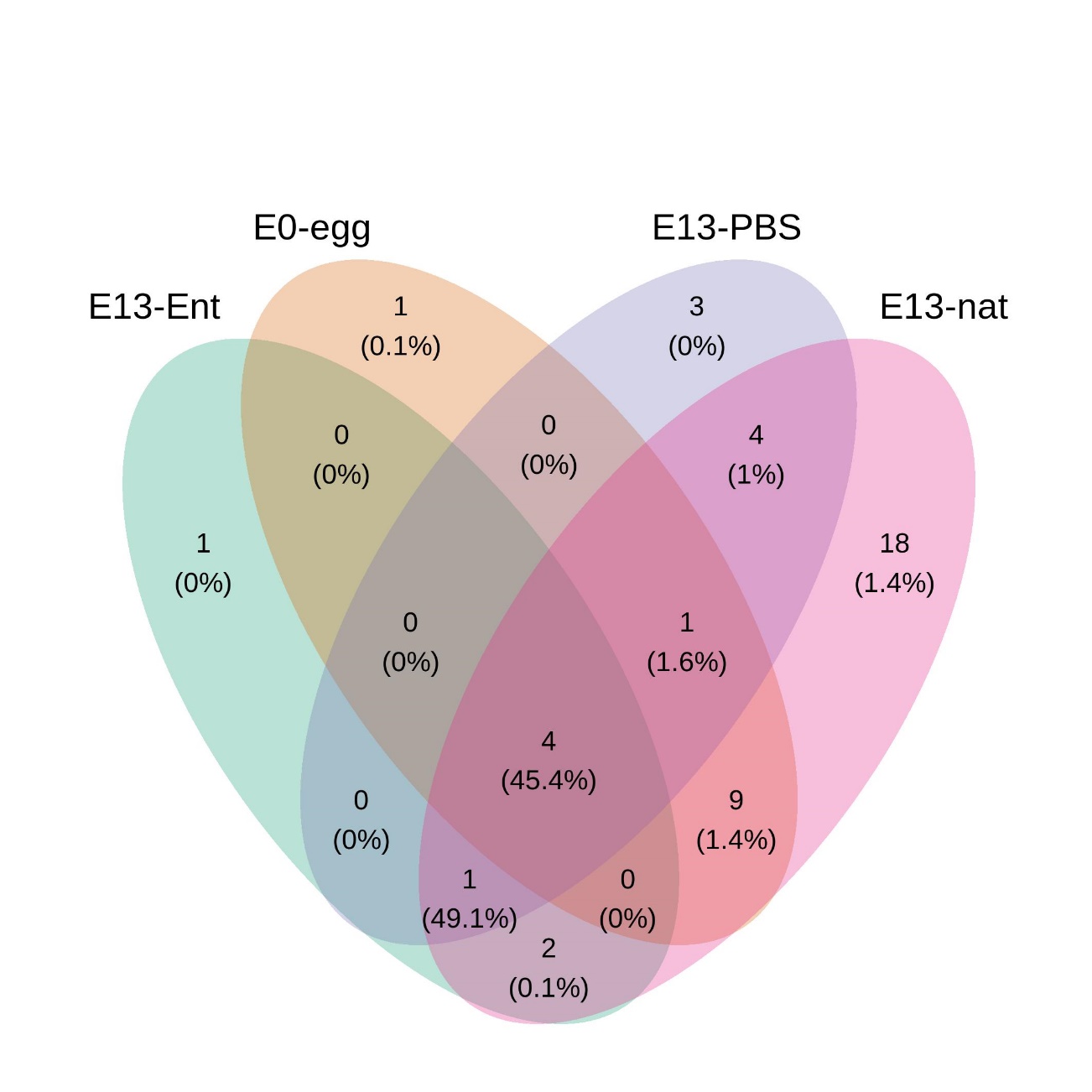

**(E)**

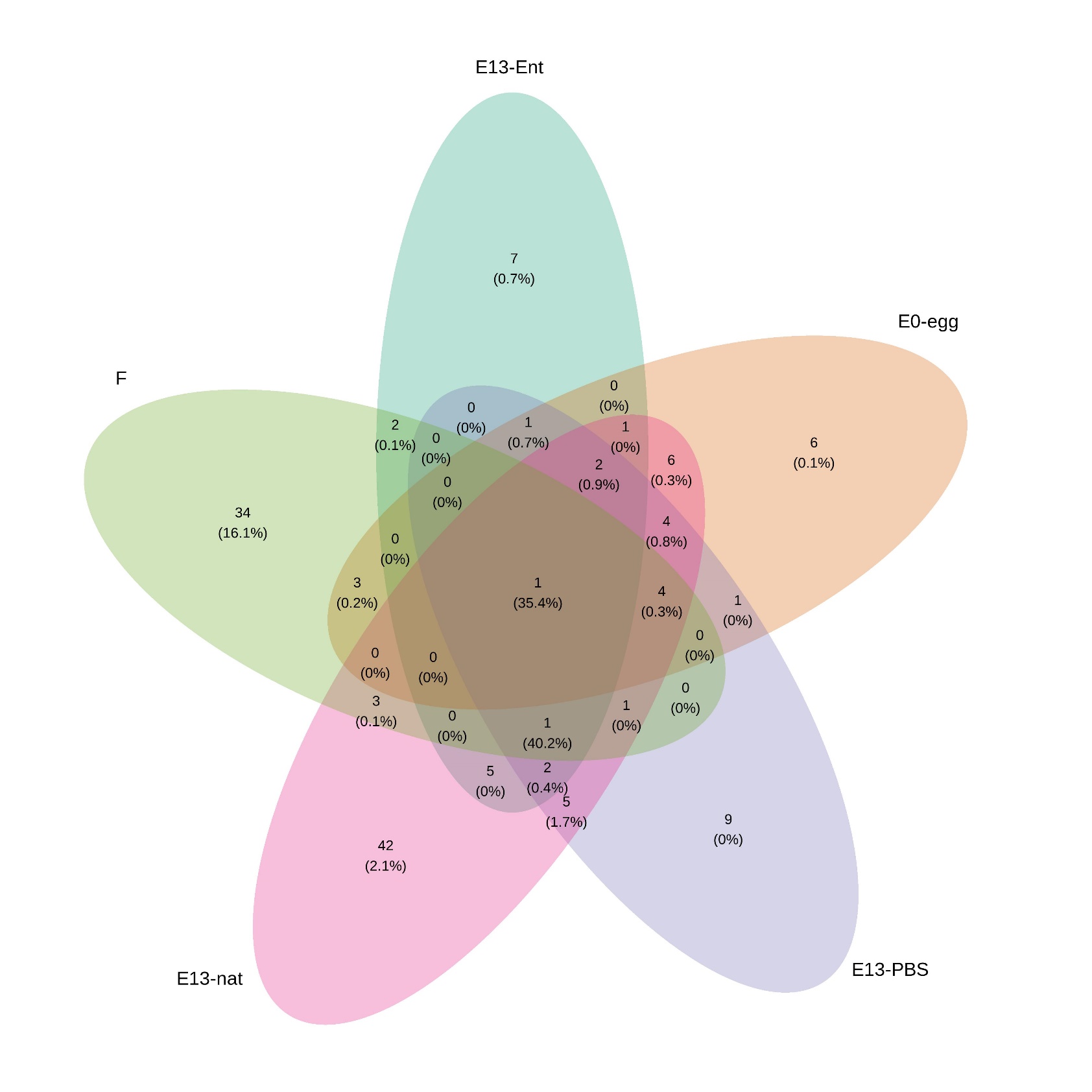

**(F)**

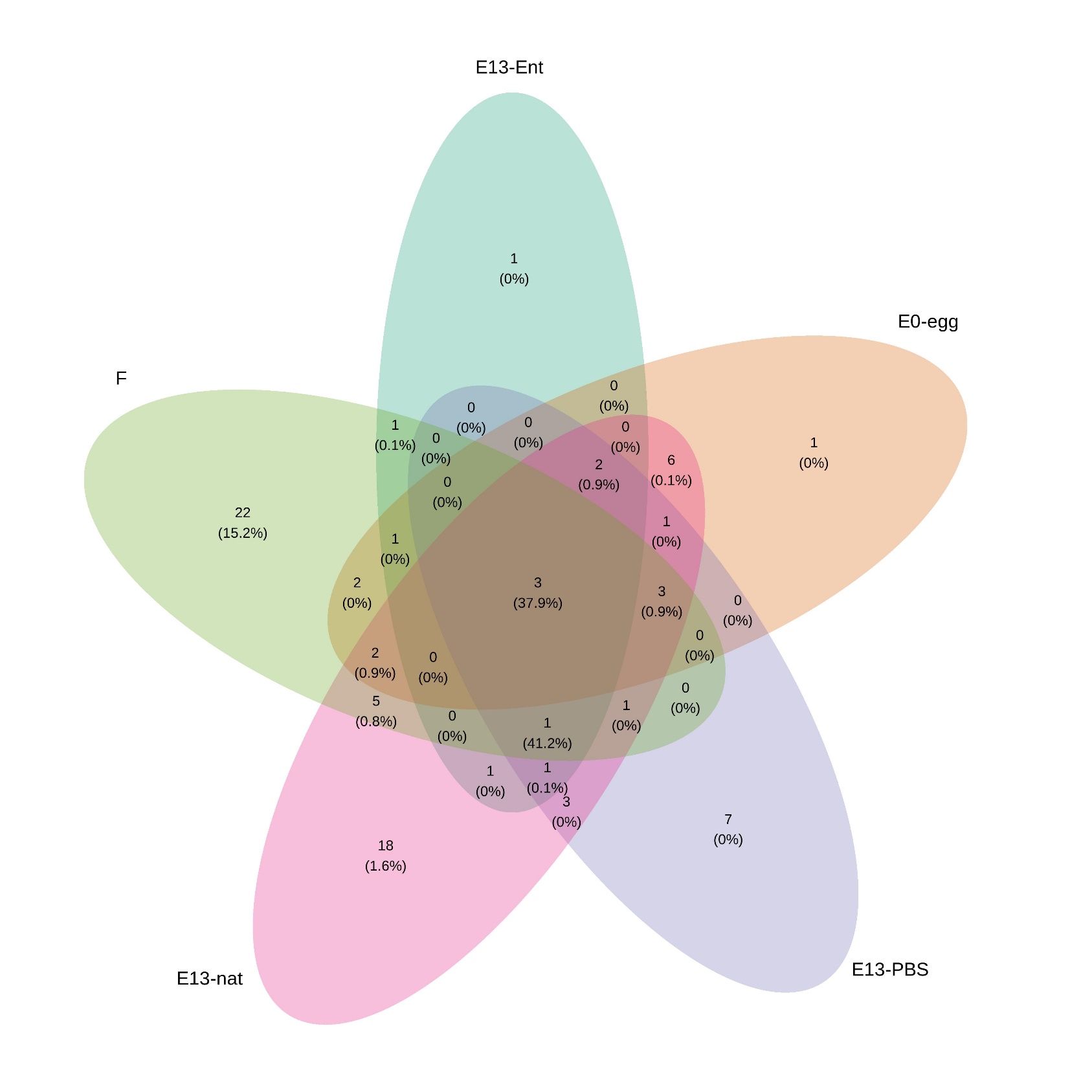

Figure S8: Predicted probability of positivity for *Corynebacterium*, *Clostridium* and *Dietzia* in egg content, embryonic intestine and female faecal samples based on the specific qPCR assays.

The qPCR and statistical analyses were always run for each combination of biological sample type and ASV separately. E0-egg – egg content sample at embryonic day 0, E13-nat – E13 intestinal sample from non-manipulated egg, F – female faecal samples, INC – negative control of isolation (i.e. isolation negative control), NTC_qPCR – negative control of PCR (i.e no template qPCR control), NTC_pre-amp – no template PCR control of preamplification step. 95 % confidence interval is shown. (A) *Dietzia* in E0-egg samples (N = 72), (B) *Dietzia* in E13-nat samples (N = 106), (C) *Clostridium* in F samples (N = 37), (D) *Corynebacterium* in F samples (N = 37) and (E) *Dietzia* in F samples (N = 42). Please note that the number of samples is given for all samples within each combination of ASV and assay (i.e. including negative controls).

(A) *Dietzia* in E0-egg samples (N = 72)

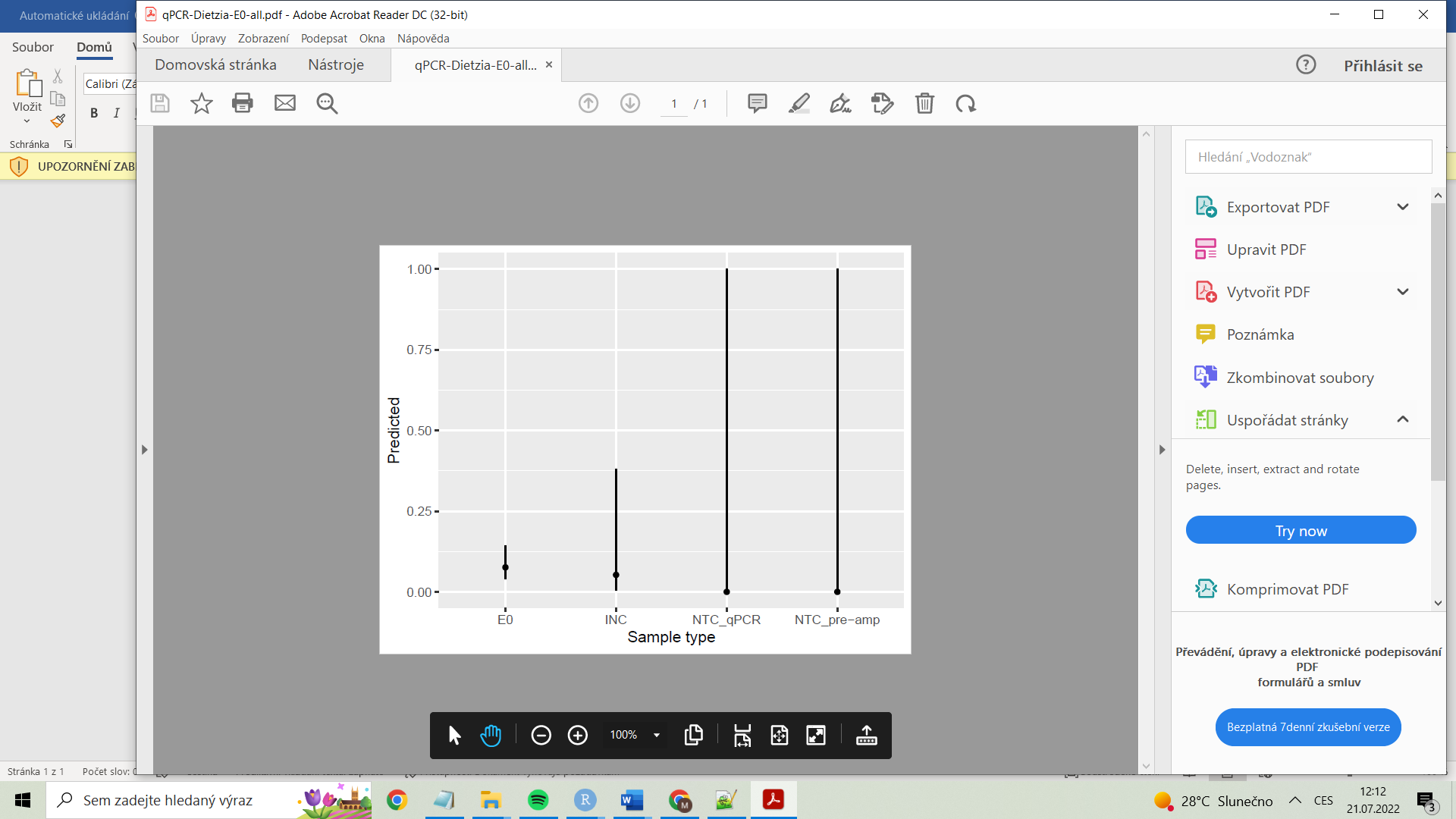

(B) *Dietzia* in E13-nat samples (N = 106)

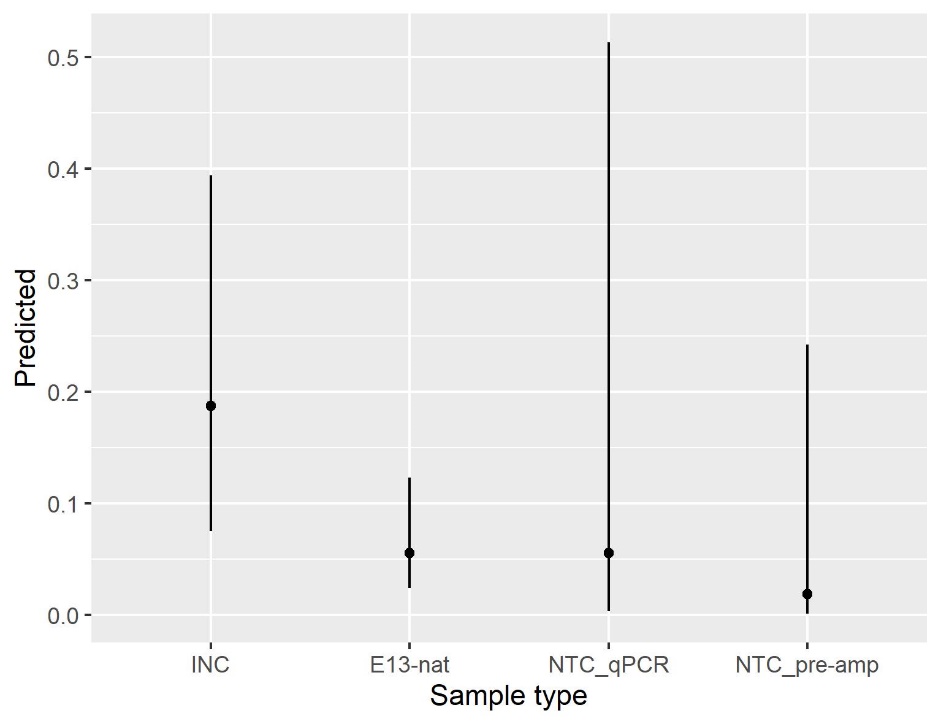

(C) *Clostridium* in all F samples (N = 37)

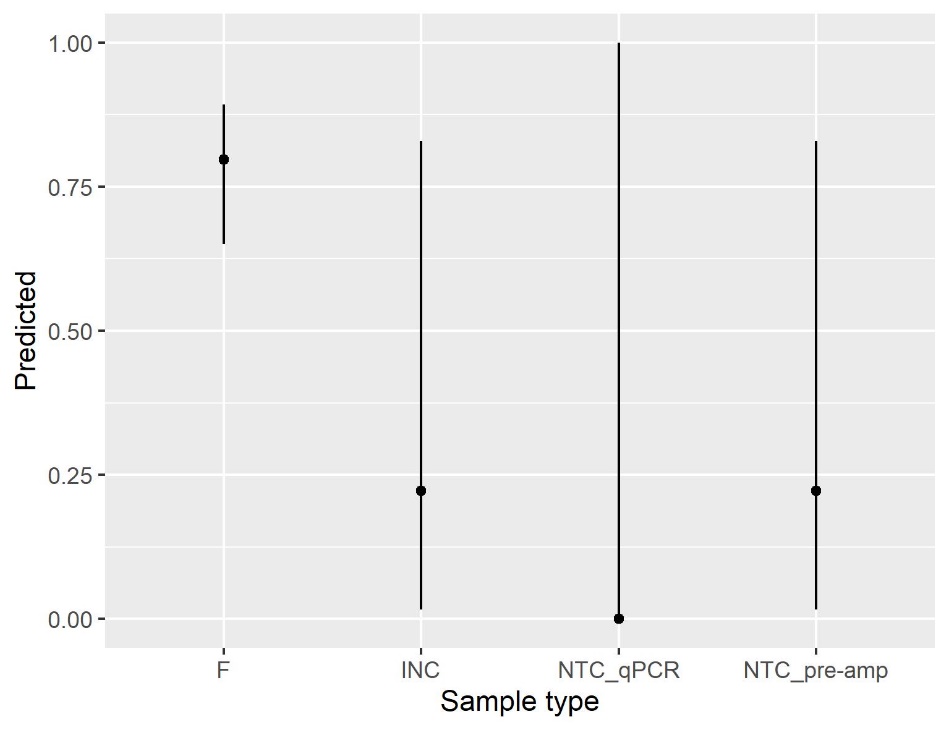

(D) *Corynebacterium* in F samples (N = 37)

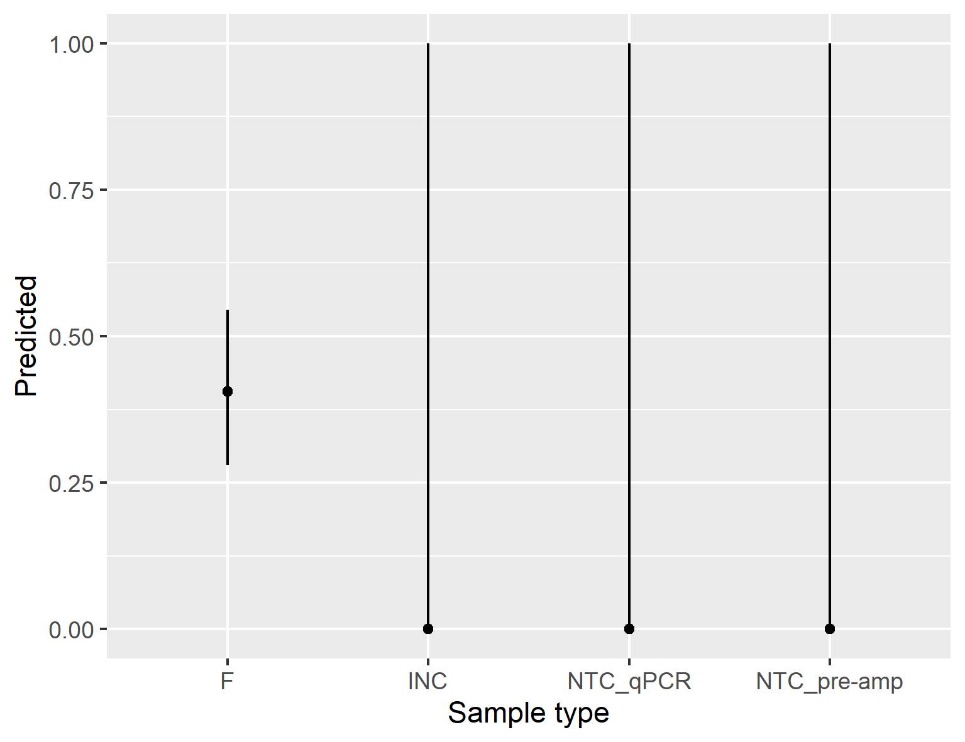

(E) *Dietzia* in all F samples (N = 42)

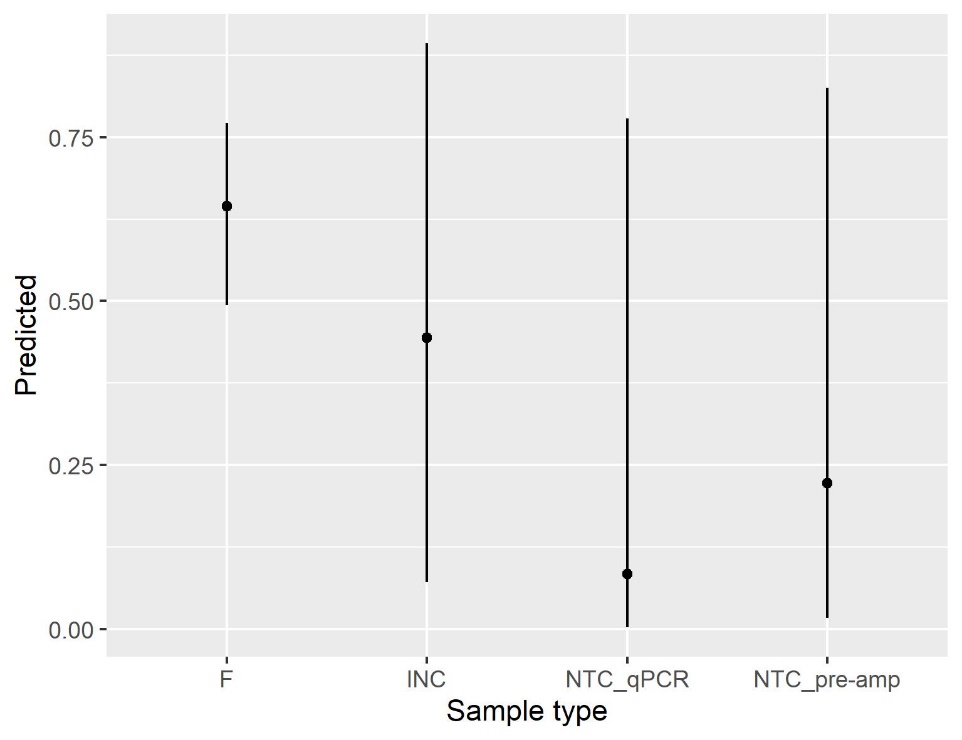

**Supplementary references**

Davis NM, Proctor DiM, Holmes SP, et al (2018) Simple statistical identification and removal of contaminant sequences in marker-gene and metagenomics data. Microbiome 6:1–14. https://doi.org/10.1186/s40168-018-0605-2

Vinkler M, Leon AE, Kirkpatrick L, et al (2018) Differing house finch cytokine expression responses to original and evolved isolates of Mycoplasma gallisepticum. Front Immunol 9:1–16. https://doi.org/10.3389/fimmu.2018.00013
