## Supplementary information 3 (SI3) for "Nearly (?) sterile avian egg in a passerine bird"

**Full model (M) details: Full output for each full model from *R* program**

Full model details are obtained for the datasets of egg content (E0-egg), embryonic gut (E13-nat) and female faecal (F) samples in the great tit. In Generalized Linear Models (GLMs), *p*-values for all terms are shown as contrasts to intercepts. The significance of each model against null model is also shown. Statistical analyses were always performed for each combination of biological sample type (i.e. E0-egg, E13-nat and F) and amplicon sequence variant separately (ASV; i.e. *Corynebacterium*, *Clostridium* and *Dietzia*), except for *Corynebacterium* and *Clostridium* in E0-egg and E13-nat, for which no samples were positive. Samples were considered positive if C_p_ < 36. The dependent variable in all models was the number of positive replicates (YES) to the number of all replicates (ALL). The explanatory variable of sample type includes either biological samples: E0-egg – egg content sample from embryonic day 0, E13-nat – E13 intestinal sample from non-manipulated egg, F – female faecal sample or different negative controls: INC – negative control of isolation (i.e. isolation negative control), NTC_qPCR – negative control of PCR (i.e. no template qPCR control), NTC_pre-amp – no template PCR control of pre-amplification step.

**List of models**

Full model details

M1: *Dietzia* in E0-egg

```
> model<-glm(cbind(YES,ALL-YES)~Sample_type,quasibinomial,data=DF)
> model2<-glm(cbind(YES,ALL-YES)~1,quasibinomial,data=DF)
> anova(model,model2,test="F")
Analysis of Deviance Table

Model 1: cbind(YES, ALL - YES) ~ Sample_type
Model 2: cbind(YES, ALL - YES) ~ 1
  Resid. Df Resid. Dev Df Deviance      F Pr(>F)
1        68     62.212
2        71     67.342 -3  -5.1304 1.1972 0.3175
> summary(model)

Call:
glm(formula = cbind(YES, ALL - YES) ~ Sample_type, family = quasibinomial,
    data = DF)

Deviance Residuals:
    Min       1Q   Median       3Q      Max
-0.6884  -0.6884  -0.6884  -0.0002   3.9327

Coefficients:
                        Estimate Std. Error t value Pr(>|t|)
(Intercept)              -2.4987     0.3589  -6.962 1.66e-09 ***
Sample_typeINC           -0.3917     1.2793  -0.306    0.760
Sample_typeNTC_pre-amp  -16.6608  2018.5249  -0.008    0.993
Sample_typeNTC_qPCR     -16.6608  3496.1876  -0.005    0.996
---
Signif. codes:  0 ‘***’ 0.001 ‘**’ 0.01 ‘*’ 0.05 ‘.’ 0.1 ‘ ’ 1

(Dispersion parameter for quasibinomial family taken to be 1.428455)

    Null deviance: 67.342  on 71  degrees of freedom
Residual deviance: 62.212  on 68  degrees of freedom
AIC: NA

Number of Fisher Scoring iterations: 17
```

M2: *Dietzia* in E13-nat

```
> model<-glm(cbind(YES,ALL-YES)~Sample_type,quasibinomial,data=DF)
> model2<-glm(cbind(YES,ALL-YES)~1,quasibinomial,data=DF)
> anova(model,model2,test="F")
Analysis of Deviance Table

Model 1: cbind(YES, ALL - YES) ~ Sample_type
Model 2: cbind(YES, ALL - YES) ~ 1
  Resid. Df Resid. Dev Df Deviance      F Pr(>F)
1       102     118.42
2       105     129.41 -3   -10.99 1.7875 0.1543
> summary(model)

Call:
glm(formula = cbind(YES, ALL - YES) ~ Sample_type, family = quasibinomial,
    data = DF)

Deviance Residuals:
    Min       1Q   Median       3Q      Max
-1.1162  -0.5856  -0.5856  -0.3349   4.1644

Coefficients:
                         Estimate Std. Error t value Pr(>|t|)
(Intercept)            -2.833e+00  4.441e-01  -6.379 5.29e-09 ***
Sample_typeINC          1.367e+00  6.910e-01   1.978   0.0506 .
Sample_typeNTC_pre-amp -1.137e+00  1.512e+00  -0.752   0.4537
Sample_typeNTC_qPCR     2.506e-15  1.539e+00   0.000   1.0000
---
Signif. codes:  0 ‘***’ 0.001 ‘**’ 0.01 ‘*’ 0.05 ‘.’ 0.1 ‘ ’ 1

(Dispersion parameter for quasibinomial family taken to be 2.049405)

    Null deviance: 129.41  on 105  degrees of freedom
Residual deviance: 118.42  on 102  degrees of freedom
AIC: NA

Number of Fisher Scoring iterations: 6
```

M3: *Clostridium* in F

```
> model<-glm(cbind(YES,ALL-YES)~Sample_type,quasibinomial,data=DF)
> model2<-glm(cbind(YES,ALL-YES)~1,quasibinomial,data=DF)
> anova(model,model2,test="F")
Analysis of Deviance Table

Model 1: cbind(YES, ALL - YES) ~ Sample_type
Model 2: cbind(YES, ALL - YES) ~ 1
  Resid. Df Resid. Dev Df Deviance      F   Pr(>F)
1        33     109.46
2        36     162.16 -3  -52.706 5.3973 0.003911 **
---
Signif. codes:  0 ‘***’ 0.001 ‘**’ 0.01 ‘*’ 0.05 ‘.’ 0.1 ‘ ’ 1
> summary(model)

Call:
glm(formula = cbind(YES, ALL - YES) ~ Sample_type, family = quasibinomial,
    data = DF)

Deviance Residuals:
    Min       1Q   Median       3Q      Max
-5.3583  -0.0005   1.1665   1.1665   2.0204

Coefficients:
                        Estimate Std. Error t value Pr(>|t|)
(Intercept)               1.3683     0.3819   3.583  0.00108 **
Sample_typeINC           -2.6210     1.4961  -1.752  0.08908 .
Sample_typeNTC_pre-amp   -2.6210     1.4961  -1.752  0.08908 .
Sample_typeNTC_qPCR     -19.6494  2946.0251  -0.007  0.99472
---
Signif. codes:  0 ‘***’ 0.001 ‘**’ 0.01 ‘*’ 0.05 ‘.’ 0.1 ‘ ’ 1

(Dispersion parameter for quasibinomial family taken to be 3.255096)

    Null deviance: 162.16  on 36  degrees of freedom
Residual deviance: 109.46  on 33  degrees of freedom
AIC: NA
```

Number of Fisher Scoring iterations: 15

M4: *Corynebacterium* in F

```
> model<-glm(cbind(YES,ALL-YES)~Sample_type,quasibinomial,data=DF)
> model2<-glm(cbind(YES,ALL-YES)~1,quasibinomial,data=DF)
> anova(model,model2,test="F")
Analysis of Deviance Table

Model 1: cbind(YES, ALL - YES) ~ Sample_type
Model 2: cbind(YES, ALL - YES) ~ 1
  Resid. Df Resid. Dev Df Deviance      F  Pr(>F)
1        33     116.56
2        36     141.59 -3  -25.028 3.0564 0.04187 *
---
Signif. codes:  0 ‘***’ 0.001 ‘**’ 0.01 ‘*’ 0.05 ‘.’ 0.1 ‘ ’ 1
> summary(model)

Call:
glm(formula = cbind(YES, ALL - YES) ~ Sample_type, family = quasibinomial,
    data = DF)

Deviance Residuals:
    Min       1Q   Median       3Q      Max
-3.0610  -1.7673  -0.2586  -0.0003   4.0292

Coefficients:
                        Estimate Std. Error t value Pr(>|t|)
(Intercept)              -0.3814     0.2864  -1.332    0.192
Sample_typeINC          -17.7781  2931.3206  -0.006    0.995
Sample_typeNTC_pre-amp  -18.6445  4520.6798  -0.004    0.997
Sample_typeNTC_qPCR     -18.6445  4520.6798  -0.004    0.997

(Dispersion parameter for quasibinomial family taken to be 2.729596)

    Null deviance: 141.59  on 36  degrees of freedom
Residual deviance: 116.56  on 33  degrees of freedom
AIC: NA

Number of Fisher Scoring iterations: 16
```

M5: *Dietzia* in F

```
> model<-glm(cbind(YES,ALL-YES)~Sample_type,quasibinomial,data=DF)
> model2<-glm(cbind(YES,ALL-YES)~1,quasibinomial,data=DF)
> anova(model,model2,test="F")
Analysis of Deviance Table

Model 1: cbind(YES, ALL - YES) ~ Sample_type
Model 2: cbind(YES, ALL - YES) ~ 1
  Resid. Df Resid. Dev Df Deviance      F Pr(>F)
1        38     154.33
2        41     175.45 -3  -21.127 2.2091 0.1028
> summary(model)

Call:
glm(formula = cbind(YES, ALL - YES) ~ Sample_type, family = quasibinomial,
    data = DF)

Deviance Residuals:
    Min       1Q   Median       3Q      Max
-4.3172  -0.3923   1.6223   1.6223   2.8098

Coefficients:
                       Estimate Std. Error t value Pr(>|t|)
(Intercept)              0.5968     0.3176   1.879   0.0679 .
Sample_typeINC          -0.8200     1.2391  -0.662   0.5121
Sample_typeNTC_pre-amp  -1.8496     1.4664  -1.261   0.2149
Sample_typeNTC_qPCR     -2.9947     1.8917  -1.583   0.1217
---
Signif. codes:  0 ‘***’ 0.001 ‘**’ 0.01 ‘*’ 0.05 ‘.’ 0.1 ‘ ’ 1

(Dispersion parameter for quasibinomial family taken to be 3.187863)

    Null deviance: 175.45  on 41  degrees of freedom
Residual deviance: 154.33  on 38  degrees of freedom
AIC: NA

Number of Fisher Scoring iterations: 5
```
